# Culture-based analysis of vaginal microbiota and its link to pregnancy outcomes: an observational cohort study in France

**DOI:** 10.1101/2024.09.11.612423

**Authors:** Laura Lesimple, Jessica Rousseau, Céline Plainvert, Luce Landraud, Nathalie Grall, François Goffinet, Pierre-Yves Ancel, Christophe Pannetier, Claire Poyart, Laurent Mandelbrot, Asmaa Tazi, The Inspire Consortium

**Affiliations:** Université Paris Cité, Inserm U1016, CNRS UMR 8104, Institut Cochin, Team Bacterial Pathogenesis and Innate Immune Signaling, Fédération Hospitalo-Universitaire Prem’Impact, Paris, France; BforCure, Montreuil, France; Université Paris Cité, CRESS, Obstetrical Perinatal and Pediatric Epidemiology Research Team, EPOPE, French Institute for Medical Research and Health INSERM, INRAE, Paris, France; Department of Bacteriology, National Reference Center for Streptococci, University Hospitals Assistance Publique – Hôpitaux de Paris, Hôpital Cochin, Fédération Hospitalo-Universitaire Prem’Impact, Paris, France; Department of Microbiology, University Hospitals Assistance Publique – Hôpitaux de Paris, Hôpital Louis Mourier, Colombes, France; Université Paris Cité, IAME INSERM U1137, Paris, France; Department of Bacteriology, University Hospitals Assistance Publique – Hôpitaux de Paris, Hôpital Bichat, Paris, France; Department of Obstetrics and Gynecology, Port-Royal Maternity, University Hospitals Paris Centre Cochin Port Royal, AP-HP, Paris, France; URC-CIC P1419, University Hospitals Paris Centre Cochin Port Royal, AP-HP, Paris, France; Department of Obstetrics and Gynecology, University Hospitals Assistance Publique – Hôpitaux de Paris, Hôpital Louis Mourier, Colombes, France

**Author notes:** Laurent Mandelbrot and Asmaa Tazi contributed equally. Corresponding author: Pr Asmaa Tazi, HUPC Cochin, Service de Bactériologie, 27 rue du Faubourg St Jacques, 75014, Paris, France., – **The authors report no conflict of interest.**.

**Keywords:** Vaginal microbiota, pregnancy outcomes, preterm birth, premature preterm rupture of membrane, preterm labor

## Abstract

**Rationale:** Increasing evidence links vaginal microbiota composition and preterm birth (PTB). However, most metagenomic studies are relatively small-sized, do not systematically adjust for confounders, and are difficult to transpose in clinical settings.

**Objective:** To identify, using routine vaginal microbiological cultures, signatures of preterm labor (PTL), preterm premature rupture of membranes (PPROM), and PTB.

**Methods:** We conducted an observational cohort study from August 2018 until June 2023 in France. Pregnant women were enrolled in three groups: control, PTL and PPROM. Demographic, clinical data, and pregnancy outcome were collected. Vaginal swabs were collected at enrollment and microbiological cultures were performed. The association between bacterial species and PTL, PPROM, and PTB was studied in univariate analyses. Adjusted odds ratio (aOR) and 95% confidence intervals (CI) were calculated in multivariable analyses adjusting for confounding variables.

**Results:** 1,848 women were included: 1,048 in the control group, 417 with PTL, and 383 with PPROM. Among women with PTL or PPROM, 328/646 (50.8%) spontaneously delivered preterm. Vaginal samples enriched in enterobacteria and in *Gardnerella* spp. were signatures of PTL. Lactobacilli depletion and enterobacteria enrichment were signatures of PPROM. In multivariable analysis, lactobacilli depletion was the strongest risk factor for spontaneous PTB (aOR 2.29, 95% CI 1.52-3.48).

**Conclusions:** Our study corroborates previous findings demonstrating the protective role of lactobacilli during pregnancy and highlights enterobacteria as signatures of PTL and PPROM. Furthermore, it provides perspectives for the management of women at high risk of PTB using standard microbiological techniques.

**Highlights:**

- Vaginal microbiological analysis of 800 women with preterm labor or PPROM
- Preterm labor signatures include enterobacteria and *Gardnerella* enrichment
- PPROM signatures include lactobacilli depletion and enterobacteria enrichment
- Lactobacilli depletion is the strongest risk factor for preterm birth

## Introduction

Preterm birth (PTB), defined as birth before 37 weeks of gestation, is a frequent and severe adverse pregnancy outcome. It concerns approx. 7% of pregnancies in industrialized countries and accounts for 14 million annual births worldwide [1]. As the leading causes of death in children under the age of five, PTB and its sequelae represent one of the most prevalent conditions in the Global Burden of Disease analysis and major worldwide health issues [2,3]. Up to 40% of spontaneous PTB (sPTB) are preceded by preterm premature rupture of membranes (PPROM), thereby increasing the risk of intrauterine infection and in utero demise [4,5]. In this context, disruption of the vaginal microbiota is increasingly considered to be a contributing factor [6–8].

The composition of the vaginal microbiota is influenced by factors such as age, hormonal fluctuations, gestational status, ethnicity, sexual behavior, and lifestyle [9–13]. Studies of vaginal microbiota, based on sequencing of vaginal samples from reproductive-age women, have identified five types of vaginal community structures designated community state types (CSTs) [11]. Four of these, designated CST-I, –II, –III, and –V, are dominated by one *Lactobacillus* species, while CST-IV is characterized by a more diverse microbiota dominated by anaerobic bacteria such as *Gardnerella* spp. and *Prevotella* spp. Although the distribution of these CSTs varies considerably according to ethnicity [14], it is generally accepted that, in reproductive-age women, a healthy vaginal microbiota is dominated by lactobacilli and is characterized by low bacterial diversity [15,16]. Conversely, a decrease in lactobacilli abundance is associated with an increase in microbiota diversity, a condition frequently referred to as vaginal dysbiosis and typically exemplified by the CST-IV. In the context of pregnancy, mounting evidence indicates a link between preterm labor (PTL), PPROM, and PTB and low-lactobacilli microbiota, characterized by high diversity and enrichment in *Gardnerella* spp. And various anaerobes [6,17–20]. In contrast, microbiota dominated by lactobacilli, particularly *Lactobacillus crispatus* (CST-I), have been associated with term birth.

Despite mounting evidence of an association between vaginal microbiota and pregnancy outcomes, integrating these observations into standard patient care practices remains challenging. Studies that have attempted to determine if pregnancy outcomes can be correlated with specific microbial signatures using conventional microbiological methods are lacking [21]. In addition, the impact of the vaginal microbiota on PTL and PPROM and their outcomes remains elusive, mainly because most studies are relatively small-sized and do not systematically adjust for confounders such as body mass index (BMI) and ethnicity [6]. Thus, the association between CST-IV or *Gardnerella* spp. and PTB appears to differ by ethnic group [22,23]. Furthermore, additional microbiota signatures, especially those of less prevalent yet pathogenic taxa such as Group B *Streptococcus* (GBS) or *Escherichia coli*, may be overlooked. Finally, the microbial factors predictive of PTB, as well as their stratification along with clinical risk factors in high-risk populations, have yet to be determined.

Our aim was to leverage conventional culture techniques on a large cohort of pregnant women to address these knowledge gaps and, systematically adjusting for confounding variables, to identify microbiota signatures of PTL and PPROM, along with microbiological risk factors for sPTB in women at high risk of PTB who were hospitalized for PTL or PPROM. A secondary objective was to investigate the relevance and potential application of routine culture-based methods for stratifying the risk of adverse pregnancy outcomes and for monitoring pregnant women in clinical practice.

## Material and Methods

### Study design

We conducted an observational, prospective, longitudinal cohort study from August 2018 until June 2023 in three maternity wards from university hospitals in the Paris area, in France. The total annual number of births at these three centers was approximately 11,000. Women were enrolled in three groups at the time of presentation: a control group and two groups at high risk of PTB, i.e., the PTL and PPROM groups (Figure 1). The eligibility criteria for maternal participation in the study included all pregnant women aged ≥ 18 years who met the following inclusion criteria: *i)* PTL: threatened PTB with intact membranes; and *ii)* PPROM: rupture of membranes before 37 weeks of gestation and before labor. The control group consisted of pregnant women who delivered at term and did not present with PTL or premature rupture of membranes. These women were enrolled at the time of GBS antenatal screening between 34 and 38 weeks of gestation, as recommended by the French guidelines for the prevention of perinatal infectious disease [24]. Women in the control group were exclusively recruited from Centers 1 and 2, whereas women in the PTL and PPROM groups were recruited from all three participating centers of the study.

**Figure 1.**
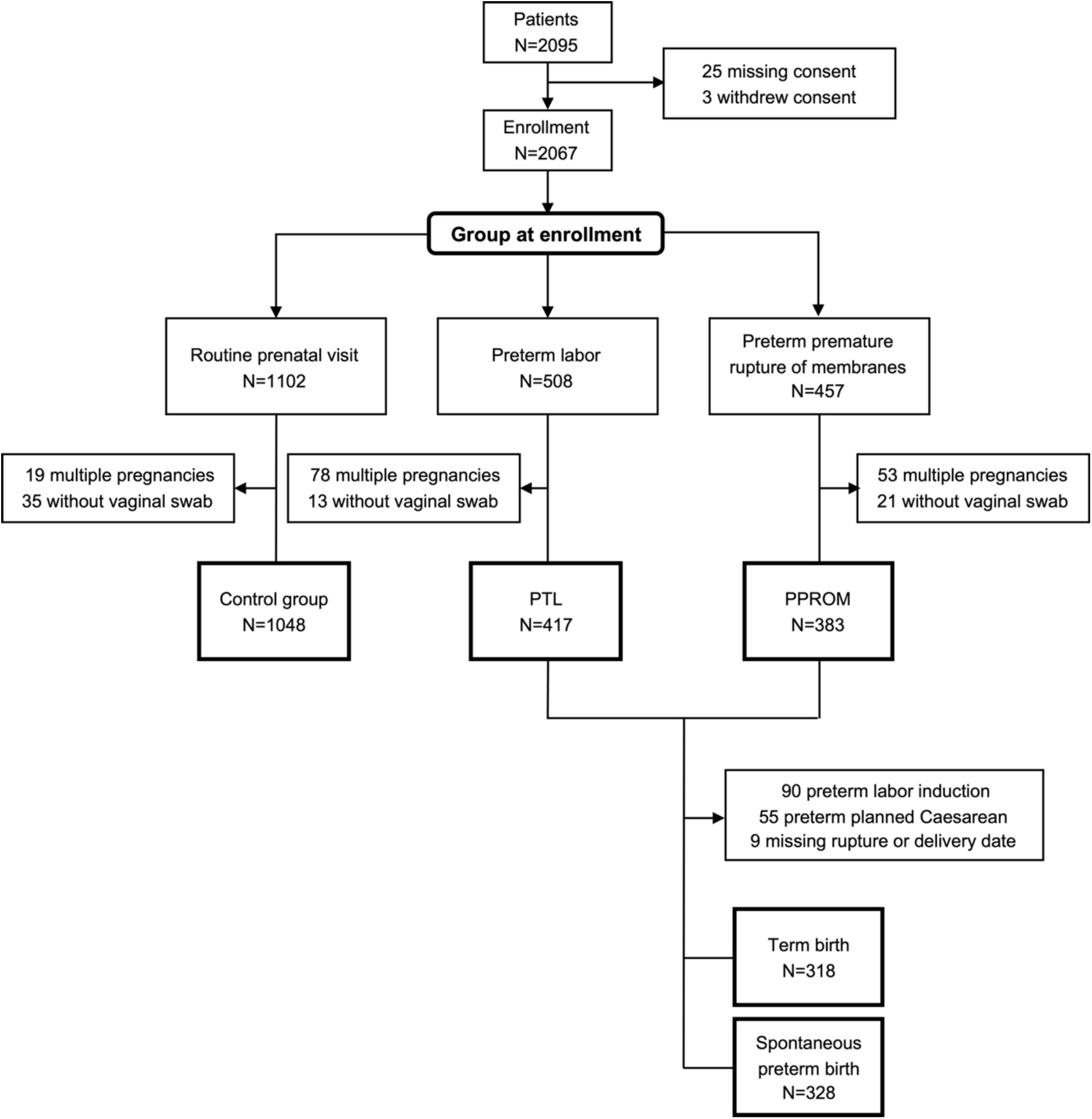
Flowchart of the study. Out of the 2,095 eligible pregnant women, 2,067 were enrolled, including 1,102 at routine prenatal visit between 34-38 weeks of gestation (control group), 508 and 457 at initial presentation for preterm labor (PTL) and preterm premature rupture of membranes (PPROM), respectively. Multiple pregnancies and women for whom a vaginal swab sample was not collected at initial presentation were excluded from the analysis, leading to cohort subgroups of 1,048 controls, 417 PTL and 383 PPROM. Women who delivered following preterm labor induction and preterm planned Caesarean section, as well as women with missing rupture or delivery date, were excluded from the analysis of the factors associated with preterm birth. The final analysis included 318 term births and 328 spontaneous preterm births. PTL: Preterm labor; PPROM: preterm premature rupture of membranes.

This study is a subset cohort study of the Innovative Strategies for Perinatal Infectious Risk Reduction (InSPIRe) project, which aimed to develop a point-of-care test to predict early-onset neonatal sepsis based on the microbiological and immunological profile of vaginal samples (ClinialTrials.gov NCT03371056). The target sample sizes for the high-risk groups were 500 women per group. The target for the control group was 1,000. The inclusions in this latter group were discontinued once this objective had been reached. The sample sizes were calculated according to the probability of early-onset neonatal sepsis in the PTL and PPROM group, i.e. 3% and 5%, respectively. A sample size of 500 women in each group was estimated sufficient to estimate a sensitivity and specificity for early-onset neonatal sepsis detection of 95% and 98% respectively. Women who did not speak or understand French and those with multiple pregnancies were excluded from this study.

The following data were collected: demographic information, clinical data, past medical and obstetrical history, and pregnancy outcome. Maternal place of birth was collected as a proxy for ethnicity, given that reporting ethnicity itself is forbidden in France.

### Sampling and laboratory procedures

Vaginal swab samples were collected at enrollment, i.e. at routine prenatal visit for the control group and upon presentation at the hospital for PTL or PPROM for the other two study groups. All vaginal swabs were obtained prior to any antibiotic administration. Vaginal swabs were sent to the laboratory in Amies transport medium (ESwab™ system, Copan Diagnostics, Italy) [25]. Specimens were immediately processed for analysis by standard routine culture methods using the following media: non-selective rich media including a Columbia agar with 5% horse blood and a chocolate agar with PolyViteX (both from bioMérieux, Marcy l’Etoile, France), incubated in aerobic atmosphere with 5% CO_2_ [26]; a UriSelect 4 chromogenic agar (Bio-Rad, Hercules, California, USA) that facilitates the detection of enterobacteria, incubated in aerobic atmosphere; a Columbia agar with 5% horse blood, nalidixic acid and colistin (bioMérieux, Marcy l’Etoile, France) for the selective isolation of anaerobes and Gram-positive cocci and a Granada agar (Fisher Scientific, Illkirch, France) for the selective isolation of GBS, both incubated under anaerobic atmosphere [26,27]. All plates were incubated at 37°C and examined after 16 and 44 hours.

All different morphotypes of bacteria and yeast colonies were identified using matrix assisted laser desorption ionization – time of flight (MALDI-TOF) mass spectrometry (Bruker Daltonics, Billerica, Massachusetts, USA) [28]. The density of vaginal colonization was estimated semi-quantitatively as follows: colonies on a single quadrant, low; colonies on more than two quadrants, high. A microbiota culture status was assigned to all vaginal samples according to the semi-quantitative assessment of the microbiological culture results. Five categories were defined (Supplementary Table 1): *Lactobacillus*-dominated, aerobic vaginitis (when dominated by a single aerobic bacterial species, including *Streptococcus* spp.), mixed vaginitis (when dominated by two or more aerobic bacterial species), anaerobes-dominated (including *Gardnerella* spp.), *Candida*-dominated, and intermediate (when the cultures yielded several bacterial genera, including lactobacilli, with no one group clearly dominating the others).

A microscopic estimation of the vaginal microbiota composition was assessed by examining the vaginal samples after Gram staining and determining the Nugent score in two out of three maternity wards according to routine practices [29]. The Nugent score, a reference method for diagnosing bacterial vaginosis, is used to determine the relative abundance of lactobacilli, small Gram-negative and Gram-variable rods (e.g., Gram-negative anaerobes and *Gardnerella* spp.) and curved Gram-negative rods (*Mobiluncus* spp.). The Nugent score ranges from 0 to 10 and is calculated by evaluating the number of morphotypes present in a microscopic field. A score ≤ 3 indicates a microbiota dominated by lactobacilli. A score comprised between 4 and 6 indicates an intermediate microbiota. A score ≥ 7 indicates a microbiota dominated by anaerobes and suggests bacterial vaginosis.

### Primary outcomes and studied factors

The primary outcome was sPTB, defined as the spontaneous onset of labor and delivery before 37 weeks of gestation. This outcome was studied in women from the PTL and PPROM groups. Individuals who delivered preterm following induction of labor or planned caesarean section were excluded from the analysis (Figure 1).

The factors studied included demographics (maternal country of birth), clinical data (BMI, gravidity, parity, tobacco use), pregnancy-related complications (gestational diabetes, hypertensive pathology, history of preterm delivery, gestational age at presentation for PTL or PPROM), and microbiological data including the microbiota culture status and the microbial species isolated from vaginal samples at enrollment. Specific attention was paid to bacteria of the *Lactobacillus* genus, and to species potentially associated with adverse outcomes including enterobacteria (such as *E. coli*, *Klebsiella* spp., *Proteus* spp.), *Staphylococcus aureus*, GBS, *Streptococcus anginosus*, *Gardnerella* spp., strict anaerobes (such as *Prevotella* spp.), and yeasts. The absence of lactobacilli in vaginal swab cultures was interpreted as “lactobacilli depletion,” whereas the presence of other species, regardless of their density, was interpreted as “enrichment.” The complete list of identified microbial species is provided in Supplementary Table 2.

### Statistical analyses

All data were collected in a computerized database and analyzed using the GraphPad Software v8.3.0 (Dotmatics, Boston, Massachusetts, USA) and RStudio v1.1.447 (Posit, Boston, Massachusetts, USA) with the following packages: R Datasets, Formal Methods and Classes, R Stats, and R Utils. Firstly, we assessed the prevalence of the studied factors according to the cohort groups as a proportion of the total number of participants in each group. Mann-Whitney tests were used to compare continuous variables, while Chi-square or Fisher’s exact tests were used to compare categorical data. Quantitative variables such as BMI and gestational age were also examined after grouping in different categories. BMI categories included underweight (<18.5), normal weight (18.5-24.9), overweight (25-29.9), and obese (≥30). Gestational age categories included extremely preterm (<28 weeks), very preterm (28-31 weeks), moderately preterm (32-36 weeks), and term (≥ 37 weeks). A *P* value < 0.05 was considered significant. Odds ratios (OR) and 95% confidence intervals (95% CI) were calculated in univariate analysis.

Secondly, to study the microbiological signatures of PTL and PPROM, and the microbiological risk factors for sPTB, multivariable logistic regression models were used with either the microbiota culture status or individual microbiological taxa as prognostic covariates. The microbiological signatures of PTL and PPROM were determined using the control group as a reference. The microbiological risk factors for sPTB were determined for women in the PTL and PPROM groups. Women in the PTL and PPROM groups who delivered at term were used as the reference group (Figure 1). All the variables of theoretical importance with a *P*-value <0.2 in the univariate analyses were included in the multivariable models, except for hypertensive disease due to the small number of affected participants. Since gravidity and parity are colinear variables, only the variable showing the most significant association in the univariate analyses was included in the multivariable analyses. Two models examined enterobacteria: one with enterobacteria as a single variable and one with the two most common species, i.e. *E. coli* and *Klebsiella pneumoniae*, as independent variables. Recruitment bias across centers –specifically, the fact that controls were recruited exclusively from Centers 1 and 2– was also accounted for in the multivariable analyses. Missing data were excluded from the analyses.

### Ethics declaration

The reported study was done in accordance with the declaration of Helsinki and approved by the French ethics committee on November, 2017 (identifier 2017 – A02755-48; NCT03371056). All individuals enrolled in the cohort provided signed informed consent that included authorization for the collection of medical information and biological samples. All collected data were anonymized and integrated in a secured electronic database.

## Results

### Study population

A total of 1,848 women were included, with 1,048 (56.7%) in the control group, 417 (22.6%) in the PTL group, and 383 (20.7%) in the PPROM group (Figure 1). The characteristics of the study population are summarized in Table 1. Among the statistically significant differences compared to the control group, the PTL and PPROM groups were characterized by a higher proportion of women born in Sub-Saharan Africa, with a history of preterm birth, and who were obese. Eventually, 31.0% (119/383) and 79.5% (209/263) of women presenting with PTL and PPROM, respectively, spontaneously delivered preterm (Table 1).

**Table 1.**
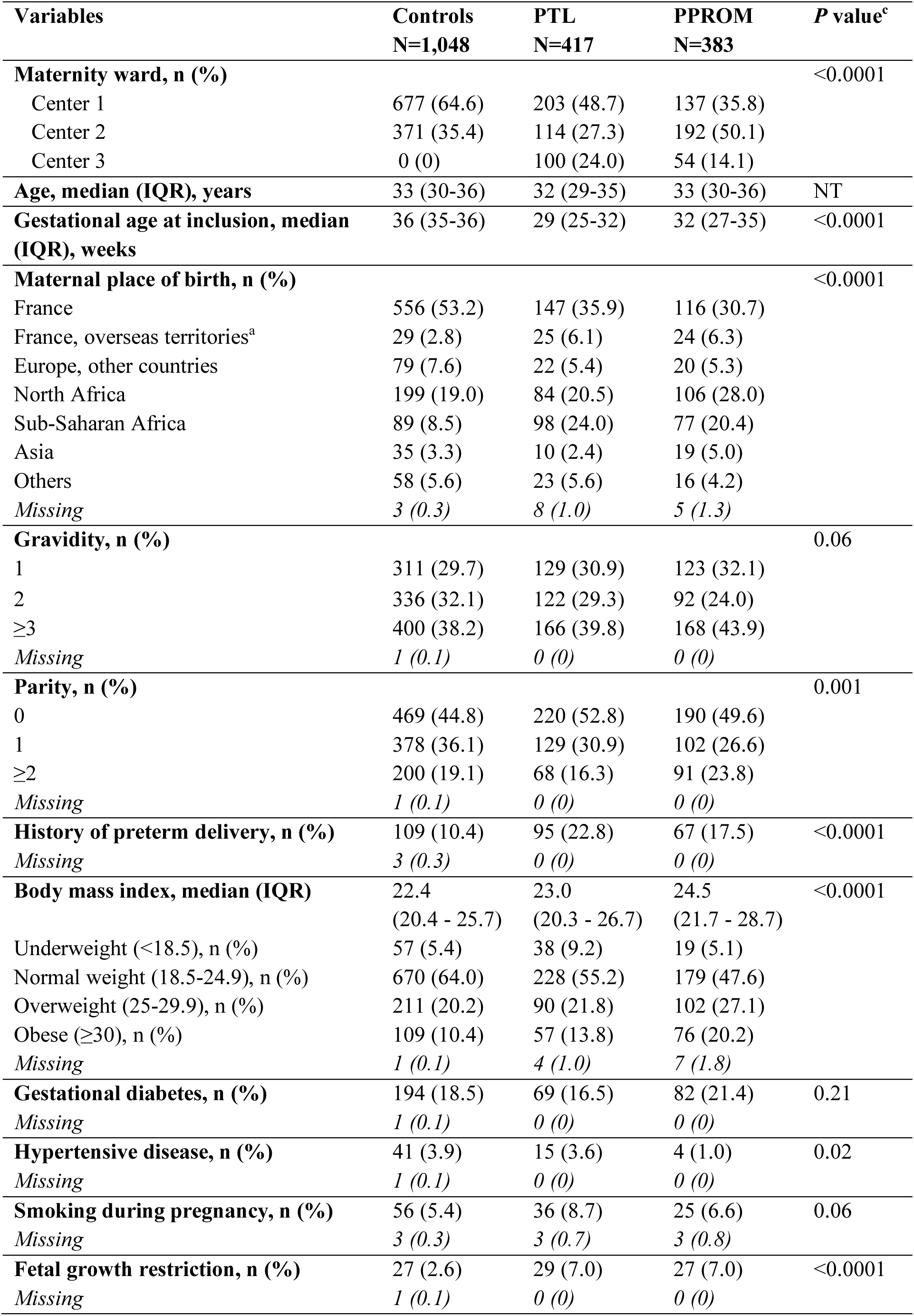
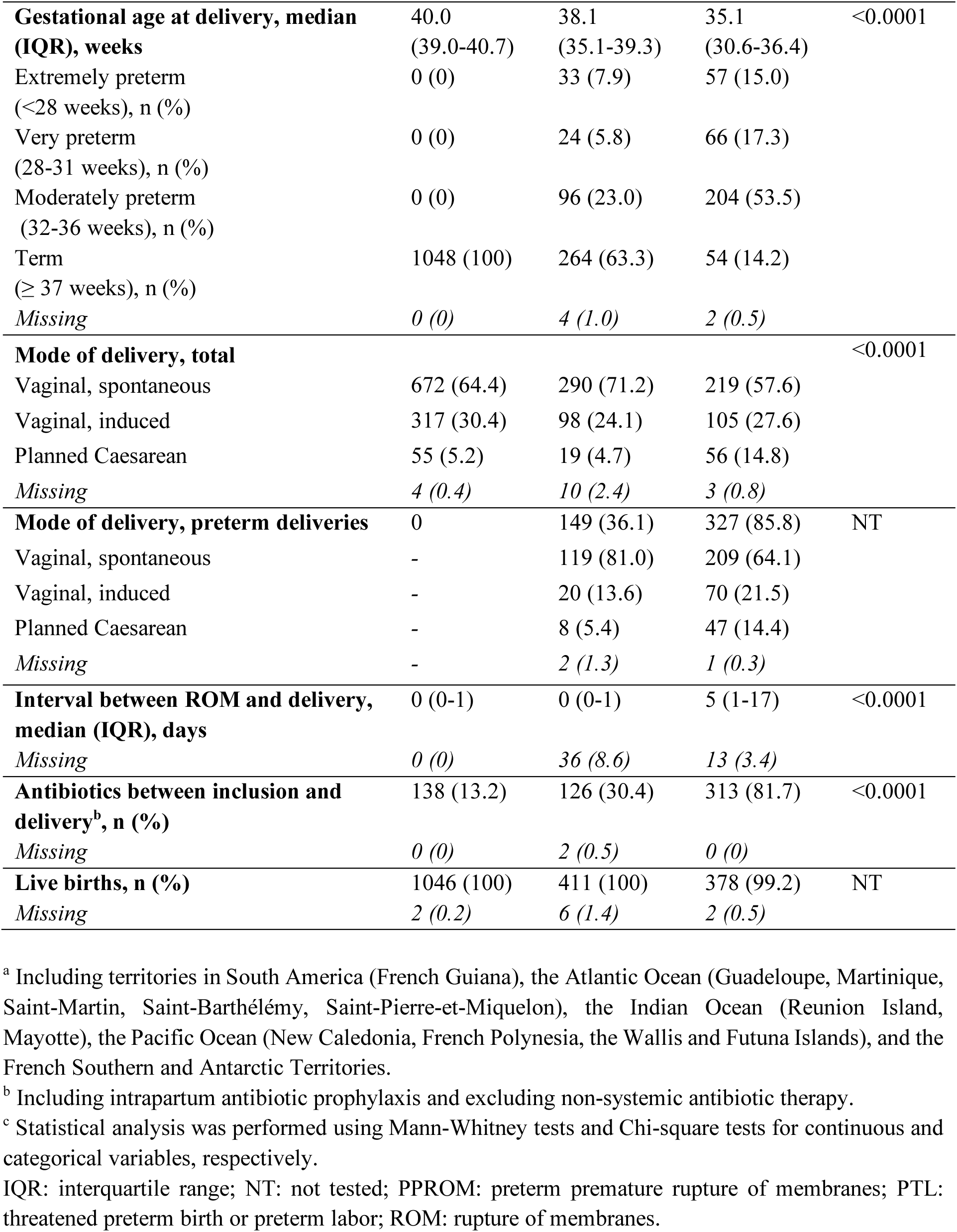
Cohort demographic and clinical characteristics.

| **Variables** | **Controls**<br>**N=1,048** | **PTL**<br>**N=417** | **PPROM**<br>**N=383** | **P value<sup>c</sup>** |
| --- | --- | --- | --- | --- |
| **Maternity ward, n (%)** |  |  |  | <0.0001 |
| Center 1 | 677 (64.6) | 203 (48.7) | 137 (35.8) |  |
| Center 2 | 371 (35.4) | 114 (27.3) | 192 (50.1) |  |
| Center 3 | 0 (0) | 100 (24.0) | 54 (14.1) |  |
| **Age, median (IQR), years** | 33 (30-36) | 32 (29-35) | 33 (30-36) | NT |
| **Gestational age at inclusion, median (IQR), weeks** | 36 (35-36) | 29 (25-32) | 32 (27-35) | <0.0001 |
| **Maternal place of birth, n (%)** |  |  |  | <0.0001 |
| France | 556 (53.2) | 147 (35.9) | 116 (30.7) |  |
| France, overseas territories<sup>a</sup> | 29 (2.8) | 25 (6.1) | 24 (6.3) |  |
| Europe, other countries | 79 (7.6) | 22 (5.4) | 20 (5.3) |  |
| North Africa | 199 (19.0) | 84 (20.5) | 106 (28.0) |  |
| Sub-Saharan Africa | 89 (8.5) | 98 (24.0) | 77 (20.4) |  |
| Asia | 35 (3.3) | 10 (2.4) | 19 (5.0) |  |
| Others | 58 (5.6) | 23 (5.6) | 16 (4.2) |  |
| Missing | 3 (0.3) | 8 (1.0) | 5 (1.3) |  |
| **Gravidity, n (%)** |  |  |  | 0.06 |
| 1 | 311 (29.7) | 129 (30.9) | 123 (32.1) |  |
| 2 | 336 (32.1) | 122 (29.3) | 92 (24.0) |  |
| ≥3 | 400 (38.2) | 166 (39.8) | 168 (43.9) |  |
| Missing | 1 (0.1) | 0 (0) | 0 (0) |  |
| **Parity, n (%)** |  |  |  | 0.001 |
| 0 | 469 (44.8) | 220 (52.8) | 190 (49.6) |  |
| 1 | 378 (36.1) | 129 (30.9) | 102 (26.6) |  |
| ≥2 | 200 (19.1) | 68 (16.3) | 91 (23.8) |  |
| Missing | 1 (0.1) | 0 (0) | 0 (0) |  |
| **History of preterm delivery, n (%)** | 109 (10.4) | 95 (22.8) | 67 (17.5) | <0.0001 |
| Missing | 3 (0.3) | 0 (0) | 0 (0) |  |
| **Body mass index, median (IQR)** | 22.4<br>(20.4 - 25.7) | 23.0<br>(20.3 - 26.7) | 24.5<br>(21.7 - 28.7) | <0.0001 |
| Underweight (<18.5), n (%) | 57 (5.4) | 38 (9.2) | 19 (5.1) |  |
| Normal weight (18.5-24.9), n (%) | 670 (64.0) | 228 (55.2) | 179 (47.6) |  |
| Overweight (25-29.9), n (%) | 211 (20.2) | 90 (21.8) | 102 (27.1) |  |
| Obese (≥30), n (%) | 109 (10.4) | 57 (13.8) | 76 (20.2) |  |
| Missing | 1 (0.1) | 4 (1.0) | 7 (1.8) |  |
| **Gestational diabetes, n (%)** | 194 (18.5) | 69 (16.5) | 82 (21.4) | 0.21 |
| Missing | 1 (0.1) | 0 (0) | 0 (0) |  |
| **Hypertensive disease, n (%)** | 41 (3.9) | 15 (3.6) | 4 (1.0) | 0.02 |
| Missing | 1 (0.1) | 0 (0) | 0 (0) |  |
| **Smoking during pregnancy, n (%)** | 56 (5.4) | 36 (8.7) | 25 (6.6) | 0.06 |
| Missing | 3 (0.3) | 3 (0.7) | 3 (0.8) |  |
| **Fetal growth restriction, n (%)** | 27 (2.6) | 29 (7.0) | 27 (7.0) | <0.0001 |
| Missing | 1 (0.1) | 0 (0) | 0 (0) |  |
| **Gestational age at delivery, median (IQR), weeks** | 40.0<br>(39.0-40.7) | 38.1<br>(35.1-39.3) | 35.1<br>(30.6-36.4) | <0.0001 |
| Extremely preterm<br>(<28 weeks), n (%) | 0 (0) | 33 (7.9) | 57 (15.0) |  |
| Very preterm<br>(28-31 weeks), n (%) | 0 (0) | 24 (5.8) | 66 (17.3) |  |
| Moderately preterm<br>(32-36 weeks), n (%) | 0 (0) | 96 (23.0) | 204 (53.5) |  |
| Term<br>(≥ 37 weeks), n (%) | 1048 (100) | 264 (63.3) | 54 (14.2) |  |
| *Missing* | 0 (0) | 4 (1.0) | 2 (0.5) |  |
| **Mode of delivery, total** |  |  |  | <0.0001 |
| Vaginal, spontaneous | 672 (64.4) | 290 (71.2) | 219 (57.6) |  |
| Vaginal, induced | 317 (30.4) | 98 (24.1) | 105 (27.6) |  |
| Planned Caesarean | 55 (5.2) | 19 (4.7) | 56 (14.8) |  |
| *Missing* | 4 (0.4) | 10 (2.4) | 3 (0.8) |  |
| **Mode of delivery, preterm deliveries** | 0 | 149 (36.1) | 327 (85.8) | NT |
| Vaginal, spontaneous | - | 119 (81.0) | 209 (64.1) |  |
| Vaginal, induced | - | 20 (13.6) | 70 (21.5) |  |
| Planned Caesarean | - | 8 (5.4) | 47 (14.4) |  |
| *Missing* | - | 2 (1.3) | 1 (0.3) |  |
| **Interval between ROM and delivery, median (IQR), days** | 0 (0-1) | 0 (0-1) | 5 (1-17) | <0.0001 |
| *Missing* | 0 (0) | 36 (8.6) | 13 (3.4) |  |
| **Antibiotics between inclusion and delivery<sup>b</sup>, n (%)** | 138 (13.2) | 126 (30.4) | 313 (81.7) | <0.0001 |
| *Missing* | 0 (0) | 2 (0.5) | 0 (0) |  |
| **Live births, n (%)** | 1046 (100) | 411 (100) | 378 (99.2) | NT |
| *Missing* | 2 (0.2) | 6 (1.4) | 2 (0.5) |  |

<sup>a</sup> Including territories in South America (French Guiana), the Atlantic Ocean (Guadeloupe, Martinique, Saint-Martin, Saint-Barthélemy, Saint-Pierre-et-Miquelon), the Indian Ocean (Reunion Island, Mayotte), the Pacific Ocean (New Caledonia, French Polynesia, the Wallis and Futuna Islands), and the French Southern and Antarctic Territories.
<sup>b</sup> Including intrapartum antibiotic prophylaxis and excluding non-systemic antibiotic therapy.
<sup>c</sup> Statistical analysis was performed using Mann-Whitney tests and Chi-square tests for continuous and categorical variables, respectively.
IQR: interquartile range; NT: not tested; PPROM: preterm premature rupture of membranes; PTL: threatened preterm birth or preterm labor; ROM: rupture of membranes.

### Vaginal microbiological signatures of PTL and PPROM

The microbiological characteristics of the vaginal samples collected at the initial hospital visit for PTL or PPROM compared to those collected from healthy controls are presented in Tables 2 and 3. These characteristics include the Nugent score, which was determined for 1,435 (77.7%) vaginal samples (Table 2), the microbiota culture status, and the culture results of the most frequently identified species (Table 3). More than 84% (1,208/1,435) of the study population had a Nugent score ≤ 3 whereas microbiological bacterial vaginosis, defined by a Nugent score ≥ 7, affected less than 5% of the women in the study, regardless of the cohort group (Table 2). Nevertheless, the distribution of the Nugent scores differed significantly between the three cohort groups. Nugent scores of 7 or higher were more prevalent in the PTL group (4.8% vs. 1.9%, *P*=0.008).

**Table 2.** Nugent scores at presentation with preterm labor (PTL) or preterm premature rupture of membranes (PPROM) compared to healthy controls.

| **Nugent score, n (%)** | **Total**<br>**N=1,848** | **Controls**<br>**N=1,048** | **PTL**<br>**N=417** | ***P* value<sup>a</sup>** | **PPROM**<br>**N=383** | ***P* value<sup>b</sup>** |
| --- | --- | --- | --- | --- | --- | --- |
| Normal (0-3) | 1,208 (84.2) | 713 (85.3) | 270 (85.7) | 0.01 | 225 (79.2) | 0.048 |
| Intermediate (4-6) | 190 (13.2) | 107 (12.8) | 30 (9.5) |  | 53 (18.7) |  |
| Bacterial vaginosis (≥7) | 37 (2.6) | 16 (1.9) | 15 (4.8) |  | 6 (2.1) |  |
| Missing | 413 (22.3) | 212 (20.2) | 102 (24.5) |  | 99 (25.8) |  |

<sup>a</sup> Statistical analysis comparing PTL to control, Chi-square test.
<sup>b</sup> Statistical analysis comparing PPROM to control, Chi-square test.
PTL: preterm labor; PPROM: preterm premature rupture of membranes.

**Table 3.** Microbiology cultures of vaginal samples at presentation with preterm labor (PTL) or preterm premature rupture of membranes (PPROM) compared to healthy controls.

| Microbiological results, n (%) | Total<br>N=1,848 | Controls<br>N=1,048 | PTL<br>N=417 | P value<sup>d</sup> | PPROM<br>N=383 | P value<sup>e</sup> |
| --- | --- | --- | --- | --- | --- | --- |
| **Microbiota culture status** |  |  |  | <0.0001 |  | <0.0001 |
| *Lactobacillus*-dominated | 1247 (68.3) | 787 (75.3) | 252 (60.6) |  | 208 (57.0) |  |
| Aerobic vaginitis<sup>a</sup> | 112 (6.1) | 36 (3.4) | 34 (8.2) |  | 42 (11.5) |  |
| - Enterobacteria, all species | 69 (3.9) | 12 (1.1) | 22 (5.3) |  | 35 (9.6) |  |
| - *Escherichia coli* | 46 (2.5) | 10 (1.0) | 15 (3.6) |  | 21 (5.8) |  |
| - *Klebsiella pneumoniae* | 13 (0.7) | 0 (0) | 5 (1.2) |  | 8 (2.2) |  |
| - Other enterobacteria | 10 (0.5) | 2 (0.2) | 2 (0.5) |  | 6 (1.6) |  |
| - Group B *Streptococcus* | 39 (2.1) | 23 (2.2) | 11 (2.6) |  | 5 (1.4) |  |
| - Others | 4 (0.2) | 1 (0.1) | 1 (0.2) |  | 2 (0.5) |  |
| Mixed vaginitis<sup>b</sup> | 35 (1.9) | 13 (1.2) | 12 (2.9) |  | 10 (2.7) |  |
| Intermediate | 297 (16.3) | 163 (15.6) | 71 (17.1) |  | 63 (17.3) |  |
| Anaerobes-dominated<sup>c</sup> | 110 (6.0) | 34 (3.3) | 40 (9.6) |  | 36 (9.9) |  |
| *Candida*-dominated | 25 (1.4) | 12 (1.2) | 7 (1.7) |  | 6 (1.6) |  |
| Missing | 22 (1.2) | 3 (0.3) | 1 (0.2) |  | 18 (4.7) |  |
| ***Lactobacillus* spp.** |  |  |  | <0.0001 |  | <0.0001 |
| Numerous | 1327 (72.6) | 845 (80.8) | 281 (67.5) |  | 201 (54.9) |  |
| Rare | 83 (4.5) | 33 (3.2) | 16 (3.8) |  | 34 (9.3) |  |
| Absent | 418 (22.9) | 168 (16.0) | 119 (28.6) |  | 131 (35.8) |  |
| Missing | 20 (1.1) | 2 (0.2) | 1 (0.2) |  | 17 (4.4) |  |
| **Enterobacteria, all species** |  |  |  | <0.0001 |  | <0.0001 |
| Numerous | 243 (13.2) | 77 (7.3) | 85 (20.4) |  | 81 (21.2) |  |
| Rare | 178 (9.6) | 91 (8.7) | 39 (9.4) |  | 48 (12.6) |  |
| Absent | 1426 (77.2) | 880 (84.0) | 293 (70.3) |  | 253 (66.2) |  |
| Missing | 1 (<0.1) | 0 (0) | 0 (0) |  | 1 (0.3) |  |
| ***Escherichia coli*** |  |  |  | <0.0001 |  | <0.0001 |
| Numerous | 162 (8.8) | 52 (5.0) | 59 (14.1) |  | 51 (13.5) |  |
| Rare | 141 (7.7) | 67 (6.4) | 33 (7.9) |  | 41 (10.8) |  |
| Absent | 1540 (83.5) | 929 (88.6) | 325 (77.9) |  | 286 (75.7) |  |
| Missing | 5 (0.3) | 0 (0) | 0 (0) |  | 5 (1.3) |  |
| ***Klebsiella pneumoniae*** |  |  |  | <0.0001 |  | <0.0001 |
| Numerous | 56 (3.0) | 13 (1.2) | 21 (5.0) |  | 22 (5.8) |  |
| Rare | 26 (1.4) | 12 (1.1) | 6 (1.4) |  | 8 (2.1) |  |
| Absent | 1761 (95.6) | 1023 (97.6) | 390 (93.5) |  | 348 (92.1) |  |
| Missing | 5 (0.3) | 0 (0) | 0 (0) |  | 5 (1.3) |  |
| ***Staphylococcus aureus*** |  |  |  | 0.07 |  | 0.64 |
| Numerous | 52 (2.8) | 24 (2.3) | 19 (4.6) |  | 9 (2.4) |  |
| Rare | 41 (2.2) | 25 (2.4) | 10 (2.4) |  | 6 (1.6) |  |
| Absent | 1735 (94.9) | 985 (95.3) | 387 (93.0) |  | 363 (96.0) |  |
| Missing | 20 (1.1) | 14 (1.3) | 1 (0.2) |  | 5 (1.3) |  |
| **Group B *Streptococcus*** |  |  |  | 0.91 |  | 0.86 |
| Numerous | 221 (12.0) | 125 (11.9) | 53 (12.7) |  | 43 (11.3) |  |
| Rare | 54 (2.9) | 32 (3.1) | 12 (2.9) |  | 10 (2.6) |  |
| Absent | 1571 (85.1) | 891 (85.0) | 352 (84.4) |  | 328 (86.1) |  |
| Missing | 2 (0.1) | 0 (0) | 0 (0) |  | 2 (0.5) |  |
| ***Streptococcus anginosus*** |  |  |  | 0.48 |  | 0.09 |
| Numerous | 58 (3.1) | 26 (2.5) | 15 (3.6) |  | 17 (4.5) |  |
| Rare | 18 (1.0) | 12 (1.1) | 4 (1.0) |  | 2 (0.5) |  |
| Absent | 1771 (95.9) | 1010 (96.4) | 398 (95.4) |  | 363 (95.0) |  |
| Missing | 1 (<0.1) | 0 (0) | 0 (0) |  | 1 (0.3) |  |
| **Anaerobes<sup>c</sup>** |  |  |  | 0.003 |  | 0.11 |
| Numerous | 238 (12.9) | 113 (10.8) | 72 (17.3) |  | 53 (13.9) |  |
| Rare | 22 (1.2) | 15 (1.4) | 5 (1.2) |  | 2 (0.5) |  |
| Absent | 1585 (85.9) | 919 (87.8) | 340 (81.5) |  | 326 (85.6) |  |
| Missing | 3 (0.2) | 1 (0.1) | 0 (0) |  | 2 (0.5) |  |
| ***Gardnerella* spp.** |  |  |  | <0.0001 |  | 0.06 |
| Numerous | 178 (9.6) | 72 (6.9) | 66 (15.8) |  | 40 (10.5) |  |
| Rare | 16 (0.9) | 10 (1.0) | 4 (1.0) |  | 2 (0.5) |  |
| Absent | 1651 (89.5) | 965 (92.2) | 347 (83.2) |  | 339 (89.0) |  |
| Missing | 3 (0.2) | 1 (0.1) | 0 (0) |  | 2 (0.5) |  |
| ***Candida* spp.** |  |  |  | 0.97 |  | 0.67 |
| Numerous | 263 (14.3) | 149 (14.3) | 61 (14.7) |  | 53 (14.0) |  |
| Rare | 51 (2.8) | 31 (3.0) | 12 (2.9) |  | 8 (2.1) |  |
| Absent | 1524 (82.9) | 865 (82.8) | 341 (82.4) |  | 318 (83.9) |  |
| Missing | 10 (0.5) | 3 (0.3) | 3 (0.7) |  | 4 (1.0) |  |

<sup>a</sup> including *Streptococcus* spp.
<sup>b</sup> Aerobic vaginitis if dominated by a single bacterial species, mixed vaginitis if dominated by two or more aerobic bacterial species.
<sup>c</sup> including *Gardnerella* spp.
<sup>d</sup> Statistical analysis comparing PTL to control, Chi-square tests.
<sup>e</sup> Statistical analysis comparing PPROM to control, Chi-square tests.
PTL: preterm labor; PPROM: preterm premature rupture of membranes.

Overall, 77.1% (1,410/1,828), 22.8% (421/1,847), 14.1% (260/1,845), and 10.5% (194/1,845) of vaginal swab cultures were positive for lactobacilli, enterobacteria, anaerobes, and *Gardnerella* spp., respectively. The microbiota culture status was correlated with the Nugent score (Supplementary Table 3, *P*<0.001) and differed significantly among the cohort groups (Table 3 and Figure 2A). Anaerobes-dominated microbiota accounted for 3.3% (34/1,045), 9.6% (40/416), and 9.9% (36/365) of the cases in the control, PTL, and PPROM groups, respectively. Additionally, aerobic vaginitis, which is not assessed by the Nugent score, was overrepresented in the PTL (8.2%, 34/416) and PPROM (11.5%, 42/365) groups compared to the control group (3.4%, 36/1,045).

**Figure 2.**
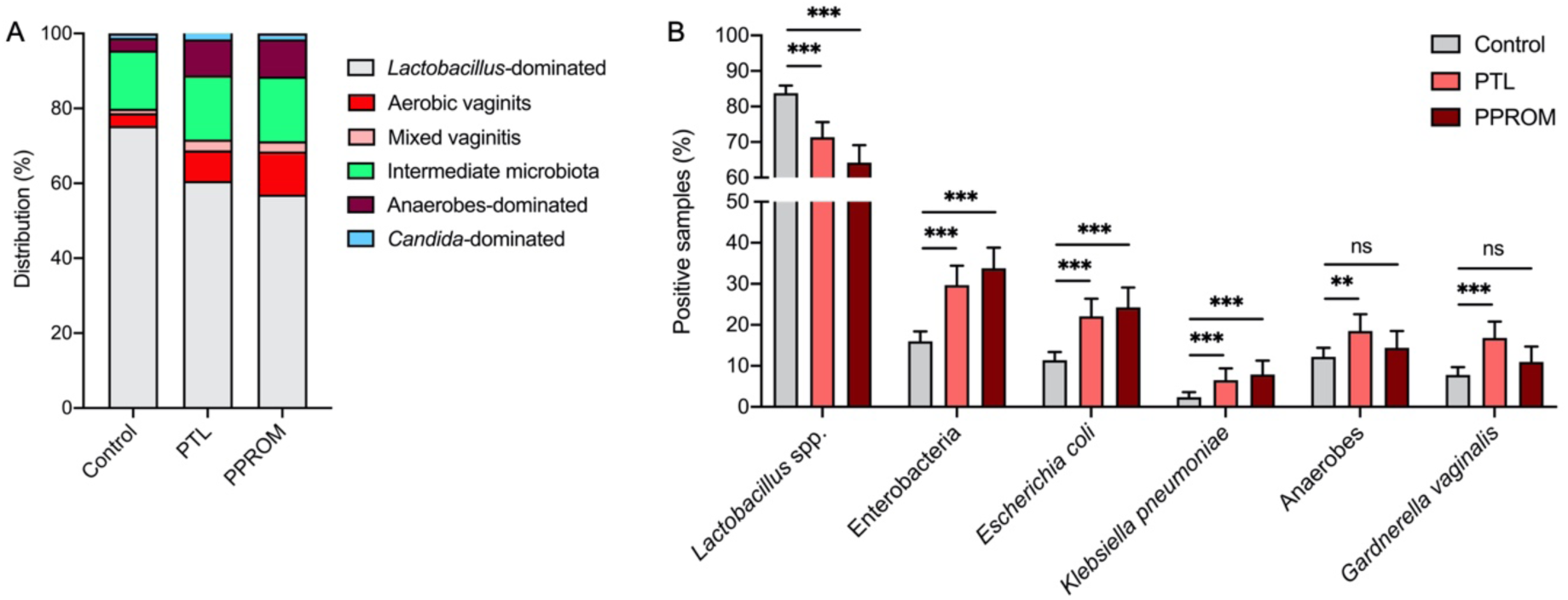
Microbiology of vaginal samples at presentation with preterm labor (PTL) or preterm rupture of membranes (PPROM). Vaginal swab cultures were compared between controls (n=1,048) and women enrolled for PTL (n=417) or PPROM (n=383). (A) Distribution of microbiota culture statuses according to the semi-quantitative assessment of the microbiological culture results. Mixed vaginitis corresponds to microbiota dominated by two or more aerobic bacterial species, including *Streptococcus* spp. Intermediate microbiota corresponds to microbiota comprising several bacterial genera, including *Lactobacillus*, with no one group clearly dominating the others. (B) Percentage of vaginal samples positive for selected bacterial group or species. (A,B) Error bars indicate the upper limit of the 95% confidence interval. ** p<0.01; *** p<0.001, ns: not significant (Chi-square tests).

The most significant differences in microbiological signatures concerned the presence and abundance of lactobacilli, enterobacteria, and anaerobes, including *Gardnerella* spp. (Table 3). Conversely, the presence and abundance of *S. aureus*, GBS and *Candida* spp. did not differ between the cohort groups. The microbial signatures of PTL included a higher proportion of samples negative for lactobacilli (28.6% vs. 16.0%, *P*<0.0002) and positive for enterobacteria (29.8% vs. 16.0%, *P*<0.0002) and anaerobes (18.5% vs. 12.2%, *P*=0.002) compared with the control group (Figure 2B). Enterobacteria were more prevalent and also more abundant (Table 3), as assessed by the number of samples with numerous bacteria compared to the total number of positive sample (68.5% in PTL *vs*. 45.8% in controls, *P*=0.0001).

The PPROM group showed a higher proportion of samples negative for lactobacilli (35.8% vs. 16.0%, *P*<0.0002) and positive for enterobacteria (33.8% vs. 16.0%, *P*<0.0002) compared with the control group. However, there was no significant difference in the proportion of samples positive for anaerobes (14.4% vs. 12.2%, *P*=0.27) (Figure 2B). Significant differences were also observed considering the abundance of lactobacilli and enterobacteria (Table 3). Lactobacilli were found in lower densities in the PPROM group (14.5% *vs*. 3.8% in controls, *P*<0.0001), whereas enterobacteria were more abundant (62.8% *vs.* 45.8% in controls, *P*=0.004).

### Multivariable analysis of microbial factors associated with PTL and PPROM

The demographic and clinical characteristics, such as country of birth, parity, and BMI, differed significantly between the cohort groups (Table 1). We investigated their association with the vaginal microbiota, as determined by Nugent scoring and microbiological cultures, in our study population. We confirmed significant variations in the Nugent scores, as well as in the presence and abundance of lactobacilli, according to country of birth (Supplementary Tables 4 and 5), gravidity and parity (Supplementary Tables 6 and 7), and BMI (Supplementary Tables 8 and 9). Notably, Nugent scores of 7 or higher and bacterial cultures negative for lactobacilli were more prevalent in women born in French overseas territories and Sub-Saharan Africa, in women with a parity ≥ 2, and in obese women. The presence and abundance of enterobacteria also varied according to the country of birth and BMI, that of anaerobes and *Gardnerella* spp. according to the country of birth, and that of GBS, *S. anginosus*, and *Candida albicans* according to parity (Supplementary Tables 5, 7 and 9).

Next, to identify the microbiological factors associated with PTL and PPROM, we performed univariate analyses (Table 4), followed by multivariable logistic regression analyses adjusted for the place of birth, parity, and BMI (Figure 3 and Supplementary Figure 1), taking the control group as the reference. In univariate analyses, the most significant risk factors for PTL were birth in French overseas territories or Sub-Saharan Africa, (OR 3.26, 95% CI 1.85-5.74, and OR 4.16, 95% CI 2.97-5.85, respectively), history of preterm delivery (OR 2.53, 95% CI 1.87-3.43), and specific microbiota culture statuses such as vaginitis due to enterobacteria and anaerobes-dominated microbiota (OR 5.73, 95% CI 2.79-11.73, and OR 3.67, 95% CI 2.28-5.93, respectively). Microbiological signatures of PTL were vaginal cultures depleted in lactobacilli and enriched in enterobacteria or *Gardnerella* spp. (OR 2.09, 95% CI 1.60-2.74, OR 2.22, 95% CI 1.70-2.90, and OR 2.37, 95% CI 1.69-3.34, respectively). In the multivariable analysis, the adjusted odds of women who presented with aerobic vaginitis due to enterobacteria and anaerobes-dominated microbiota experiencing PTL were 4.89 (95% CI 2.14-11.59) and 2.55 (95% CI 1.34-4.78) times higher, respectively, than for those with *Lactobacillus*-dominated microbiota (Figure 3A). Those of women with vaginal microbiological cultures enriched in enterobacteria or *Gardnerella* spp. were 2.67 (95% CI 1.91-3.74), and 3.88 (95% CI 1.52-11.47) times higher, respectively, than for those without such cultures (Figure 3B).

**Figure 3.**
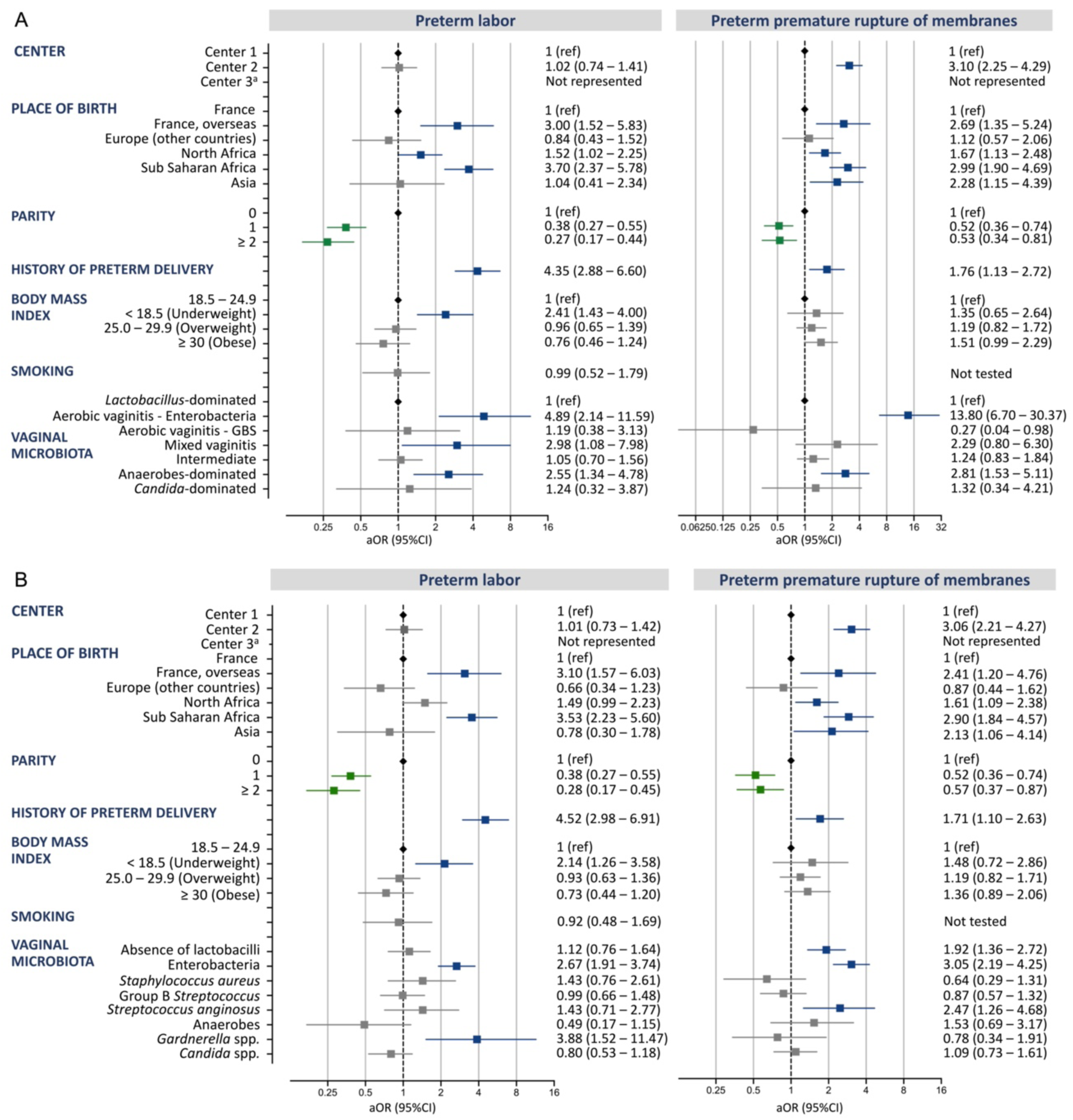
Multivariable analysis of risk factors for preterm labor and preterm premature rupture of membranes. (A) Multivariable analysis with microbiota culture status as a covariate.^a^ aOR (95% CI) of Center 3 for preterm labor and preterm premature rupture of membranes: 1.42×10^8^ (6.10×10^2^-1.03×10^70^) and 1.81×10^8^ (0.07-1.70×10^85^), respectively. (B) Multivariable analysis with microbiological taxa as covariates.^a^ aOR (95% CI) of Center 3 for preterm labor and preterm premature rupture of membranes: 1.7×10^8^ (7.3×10^2^-2.2×10^70^) and 2.4×10^8^ (0.11-1.0×10^85^), respectively. (A,B) Error bars indicate the upper and lower limit of the 95% confidence interval (CI); aOR: adjusted odds ratio; GBS: Group B *Streptococcus*.

**Table 4.**
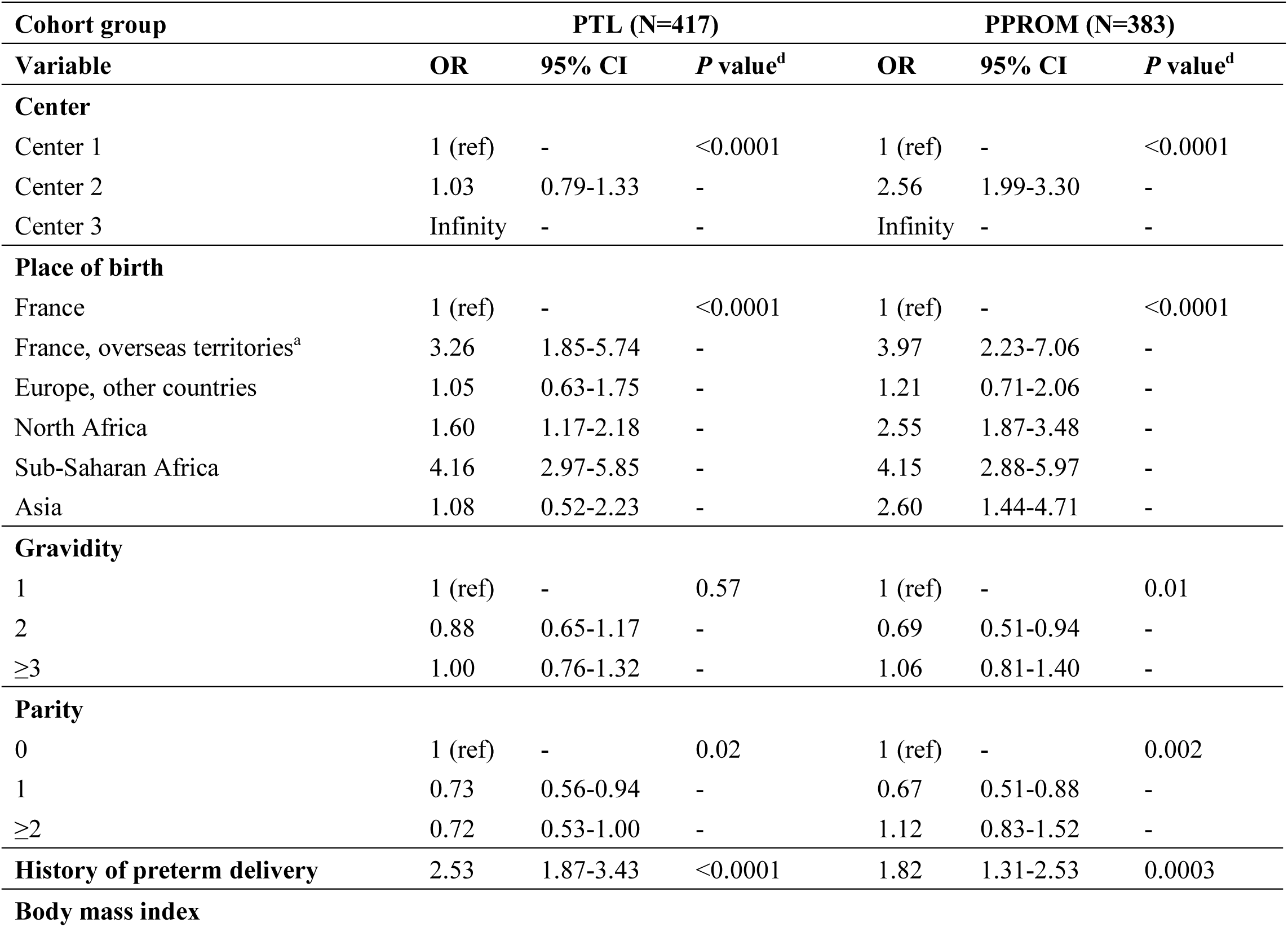
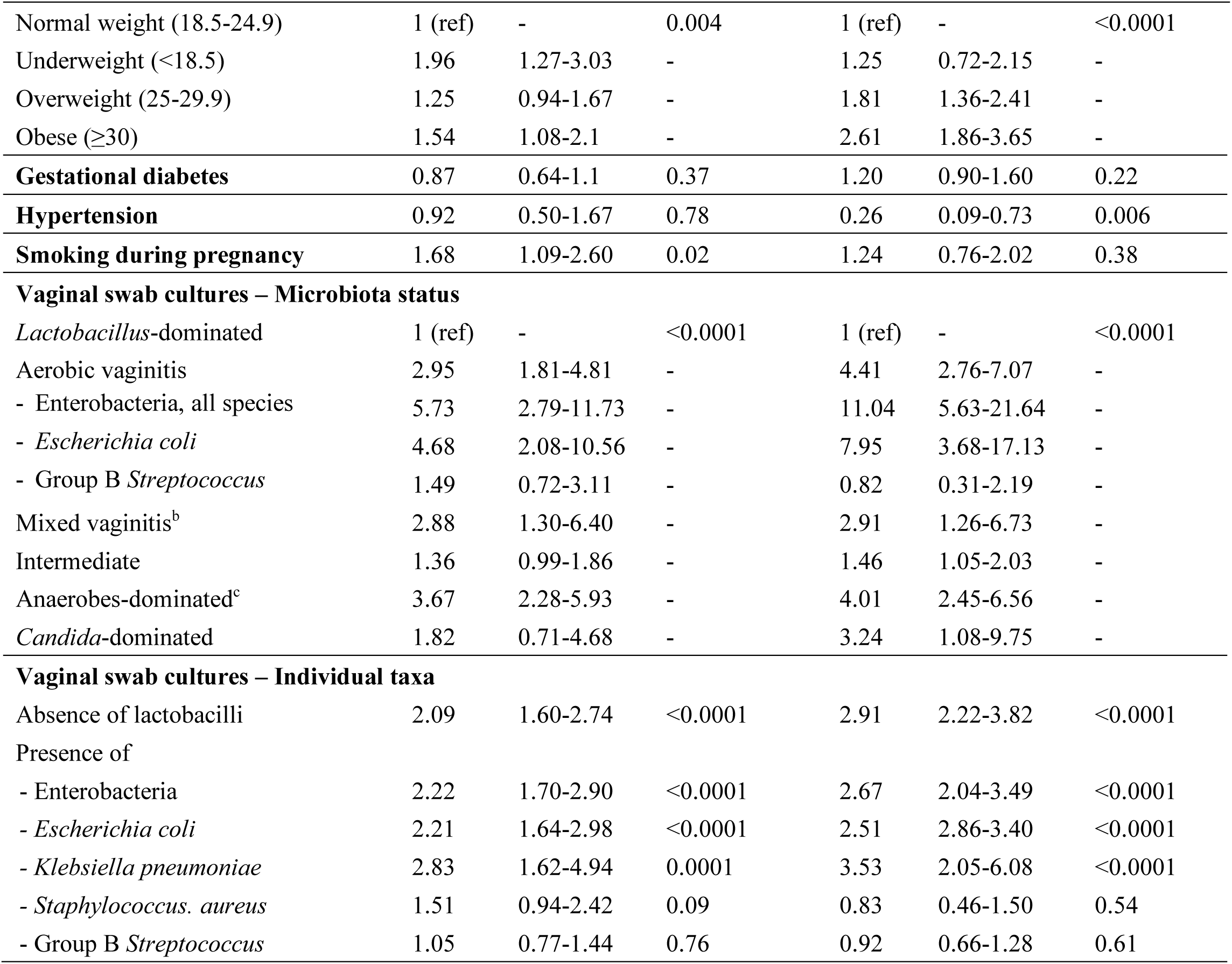
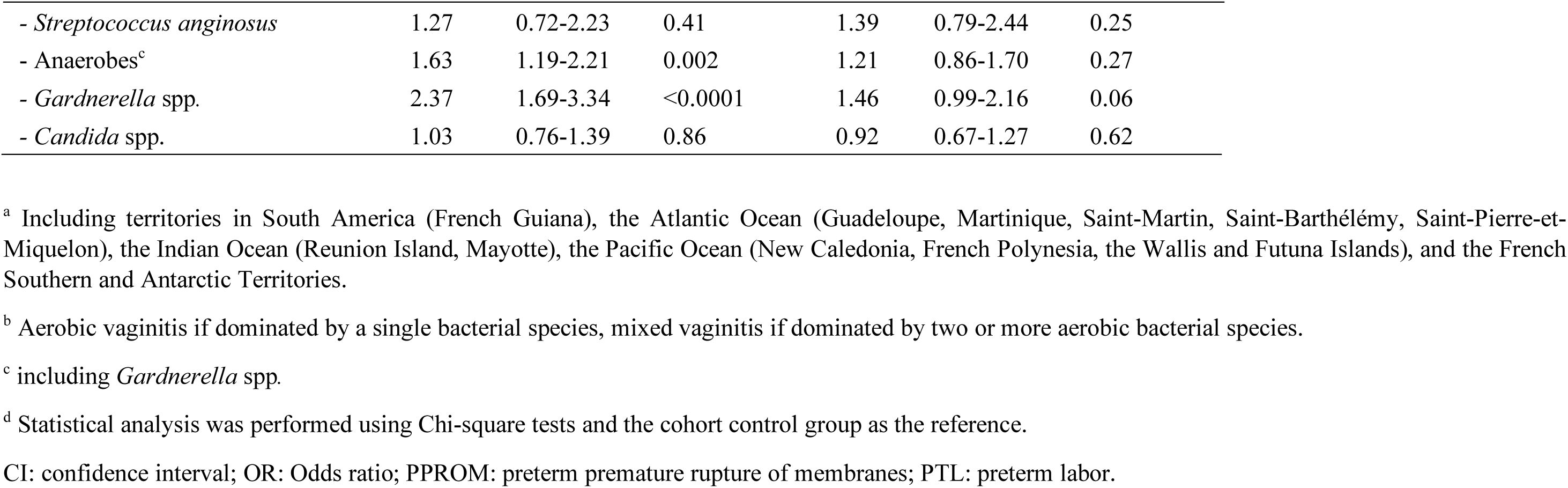
Univariate analysis of factors associated with preterm labor (PTL) and preterm premature rupture of membranes (PPROM).

**Table 5.**
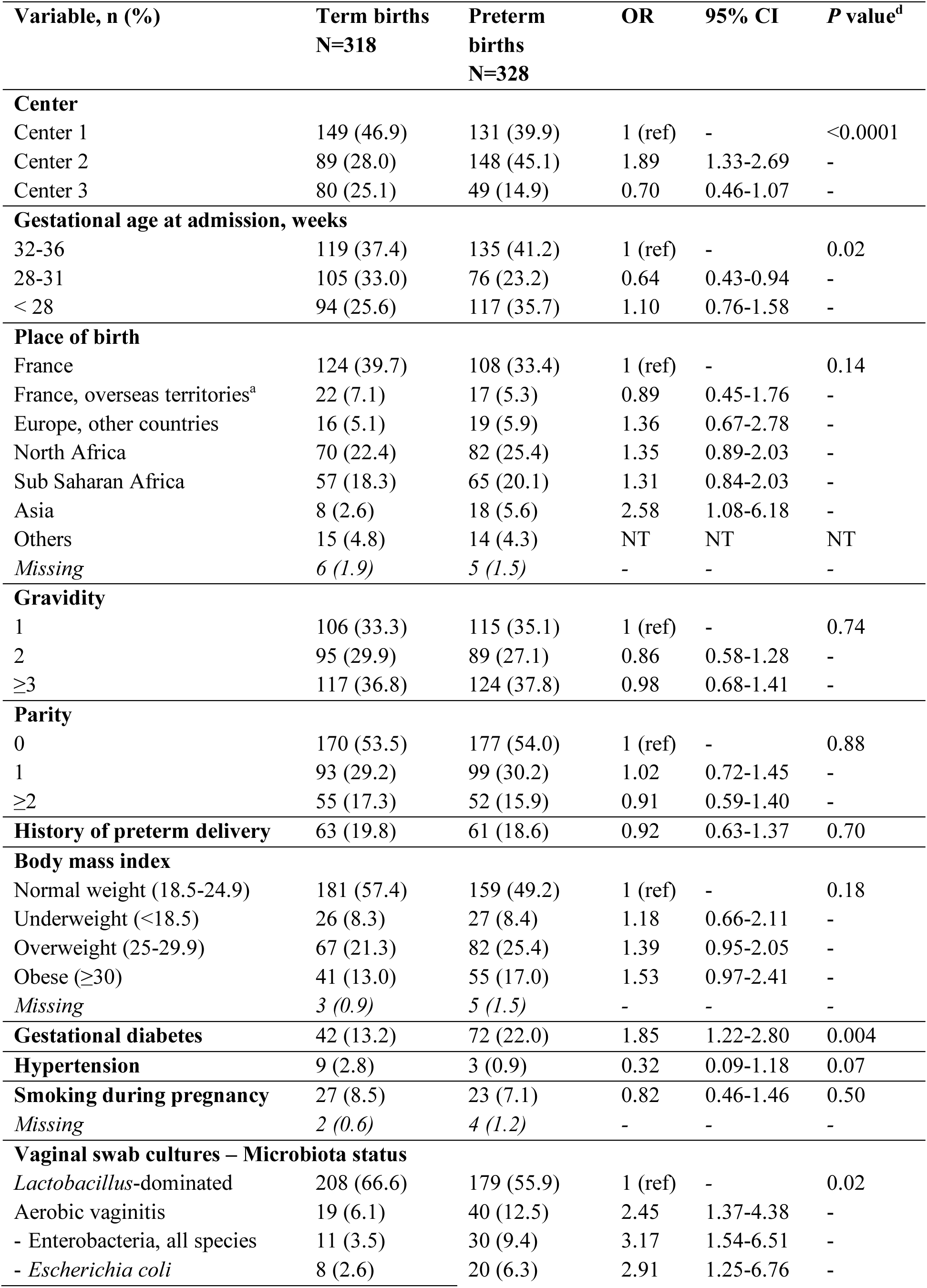
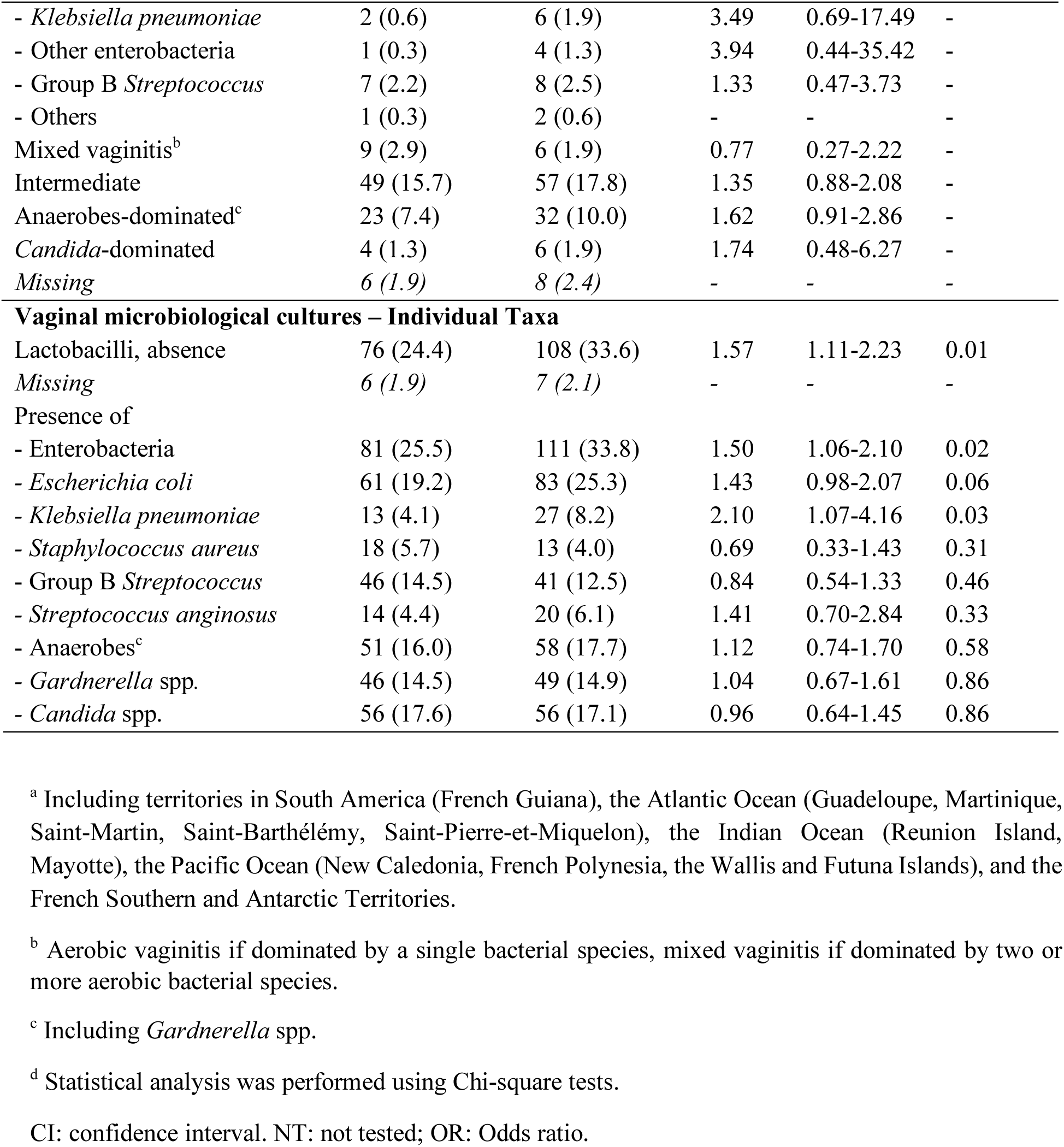
Risk factors for spontaneous preterm birth.

In the context of PPROM, in univariate analyses, the most significant risk factors were maternal birth outside Europe, with respective OR of 3.97 (95% CI 2.23-7.06) for French overseas territories, 2.55 (95% CI 1.87-3.48) for North Africa, 4.15 (95% CI 2.88-5.97) for Sub-Saharan Africa, and 2.60 (95% CI 1.44-4.71) for Asia, as well as obesity (OR 2.61, 95% CI 1.86-3.65), and non-*Lactobacillus*-dominated microbiota, with the exception of aerobic vaginitis due to GBS (Table 4). Microbiological signatures of PPROM included vaginal cultures depleted in lactobacilli (OR 2.91, 95% CI 2.22-3.82) or enriched in enterobacteria (OR 2.67, 95% CI 2.04-3.49). In the multivariable analysis, the adjusted odds of women who presented with aerobic vaginitis due to enterobacteria and anaerobes-dominated microbiota experiencing PPROM were 13.80 (95% CI 6.70-30.37) and 2.81 (95% CI 1.53-5.11) times higher, respectively, than for those with *Lactobacillus*-dominated microbiota (Figure 3A). Those of women with vaginal microbiological cultures depleted in lactobacilli and enriched in enterobacteria were 1.92 (95% CI 1.36-2.72) and 3.05 (95% CI 2.19-4.25) times higher, respectively, than for those without such cultures (Figure 3B). In addition, vaginal cultures positive for *S. anginosus* were also associated with PPROM (aOR 2.47, 95% CI 1.26-4.68).

Overall, the presence of enterobacteria in vaginal swab cultures increased the odds of PTL and PPROM, whereas the presence of *Gardnerella* spp. and the absence of lactobacilli were risk factors for PTL and PPROM, respectively. Similar results were obtained with the second model of multivariable logistic regression including *E. coli* and *K. pneumoniae* instead of enterobacteria (Supplementary Figure 1), showing that vaginal colonization by *E. coli* and *K. pneumoniae* increased the odds of PTL (aOR 2.54, 95% CI 1.75-3.69 and 2.34, 95% CI 1.17-4.65, respectively) and PPROM (aOR 2.77, 95% CI 1.91-3.99 and 2.79, 95% CI 1.47-5.34, respectively).

### Depletion in vaginal lactobacilli represents the strongest risk factor for spontaneous preterm birth

Finally, we sought to determine whether the vaginal microbiota composition at the initial hospital visit for PTL or PPROM was associated with an increased risk of sPTB. To address this question, we compared the demographic, clinical, and microbiological characteristics of the vaginal samples collected from women in the PTL and PPROM groups at enrollment who went on to deliver preterm spontaneously, with those who delivered at term (Figure 1). In univariate analyses, we found that, in addition to expected clinical factors such as gestational diabetes (OR 1.85, 95% CI 1.22-2.80), women who presented with aerobic vaginitis due to enterobacteria and vaginal microbiological cultures depleted in lactobacilli and enriched in enterobacteria were at higher risk of spontaneous preterm delivery (OR 3.17, 95% CI 1.54-6.51, OR 1.57, 95% CI 1.11-2.23, and OR 1.50, 95% CI 1.06-2.10, respectively) (Table 5).

In the multivariable analysis using microbiota culture status as a covariate, aerobic vaginitis due to enterobacteria, intermediate microbiota, and anaerobes-dominated microbiota were risk factors for sPTB (aOR 3.64, 95% CI 1.75-8.07, aOR 1.67, 95% CI 1.03-2.71, and aOR 2.18, 95% CI 1.17-4.14, respectively) (Figure 4A). In the multivariable model analyzing individual microbiological signatures, the absence of lactobacilli was the strongest risk factor for sPTB, with an aOR of 2.29 (95% CI 1.52-3.48) (Figure 4B). Lastly, a model of multivariable logistic regression where the vaginal colonization by *E. coli* and *K. pneumoniae* was analyzed showed similar results, the absence of lactobacilli being associated with sPTB (aOR 2.29, 95% CI 1.52-3.48) (Supplementary Figure 2). Additionally, the presence of *K. pneumoniae* in vaginal samples exhibited a tendency, although not statistically significant, to be linked with sPTB (aOR 1.86, 95% CI 0.90-3.99, *P* value=0.10).

**Figure 4.**
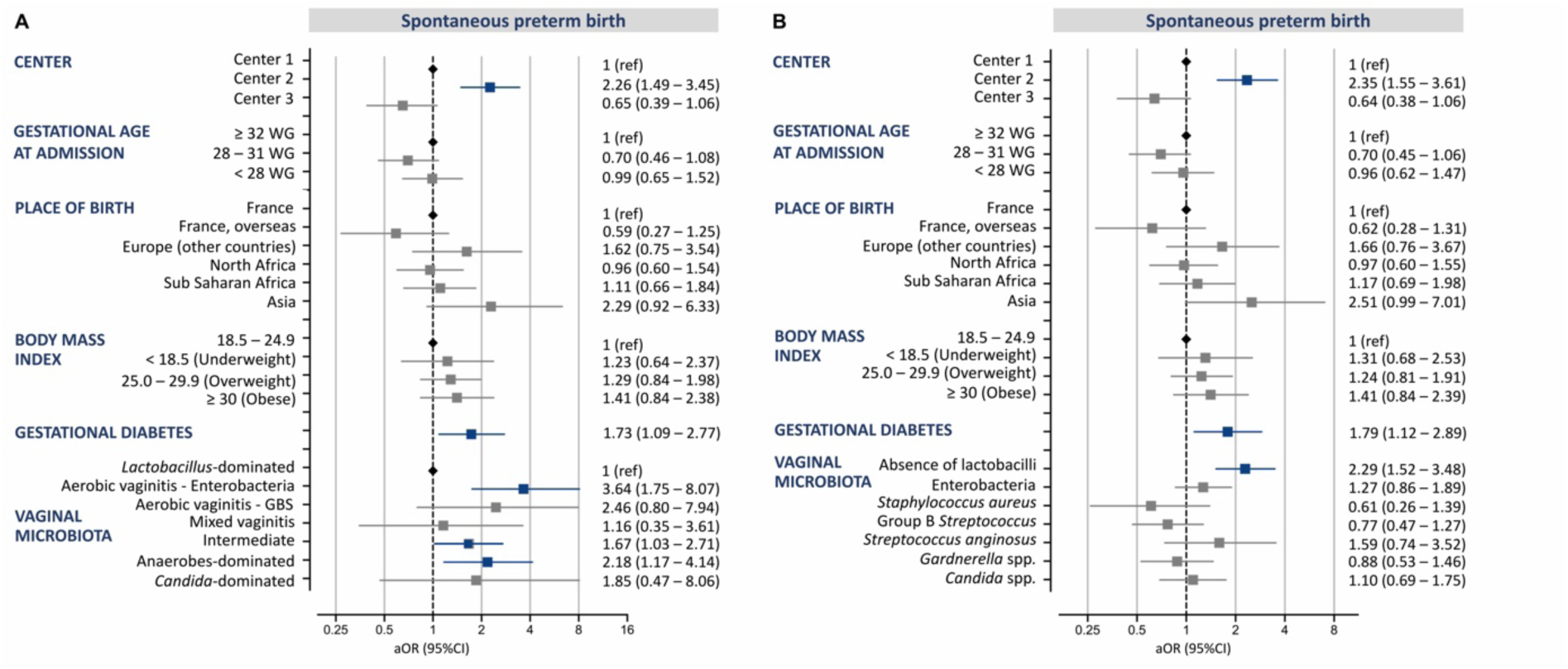
Multivariable analysis of risk factors for spontaneous preterm birth. (A) Multivariable analysis with microbiota culture status as a covariate. (B) Multivariable analysis with microbiological taxa as covariates. Error bars indicate the upper and lower limit of the 95% confidence interval (CI); aOR: adjusted odds ratio; GBS: Group B *Streptococcus*; WG: weeks of gestation.

## Discussion

### Principal Findings

In this prospective cohort study of over 1,800 pregnant women, we identified, using simple microbiological cultures of vaginal samples, distinct microbiological signatures of PTL and PPROM. Vaginal microbiological cultures suggestive of aerobic vaginitis due to enterobacteria and anaerobes-dominated microbiota were associated with PTL, PPROM, and sPTB. In addition, vaginal samples positive for enterobacteria and *Gardnerella* spp. were overrepresented in women experiencing PTL, and those positive for enterobacteria and depleted in lactobacilli were overrepresented during PPROM. Importantly, the composition of the vaginal microbiota at the initial hospital visit for PTL or PPROM correlated with the risk of sPTB, with increased odds in women with *Lactobacillus*-depleted vaginal samples.

### Results in the Context of What is Known

Our results support previous findings regarding the importance of a rich *Lactobacillus* microbiota for pregnancy outcome. Reduced *Lactobacillus* spp. abundance and increased vaginal bacterial diversity were identified as risk factors for PPROM in a study that analyzed the vaginal microbiome of 86 women [30]. A meta-analysis conducted by Gudnadottir *et al.* in which the vaginal microbiome was assessed before the onset of labor reported an increased risk of preterm delivery in women with low-lactobacilli microbiota [18]. Lastly, in a recent longitudinal cohort study of 175 pregnant women comprising 40 PTB, the vaginal microbiome of pregnancies resulting in PTB was found to be genetically more diverse, a diversity which was driven by *Gardnerella* spp. [23]. In our study, *Gardnerella* spp. were more prevalent in the vaginal samples of women presenting with PTL. However, it was not identified *per se* as a risk factor for PTB, unlike anaerobes-dominated microbiota. This discrepancy is likely attributable to the study design, as it focused exclusively on the risk factors for sPTB in women already experiencing PTL or PPROM. Furthermore, the substantial size of our cohort study elucidated the previously unrecognized risk associated with enterobacteria in PPROM and PTL, a finding that had been overlooked in smaller studies. Similar findings were demonstrated in a subset of our cohort study using shotgun metagenomics on samples collected at delivery, in which *E. coli* and *K. pneumoniae* were identified as primary drivers of PTL and PPROM [19].

### Clinical Implications

The role of bacterial vaginosis and *Gardnerella* spp. in PTL and PTB has long been suspected [31,32]. Increasing evidence now supports a direct role for *G. vaginalis* in the pathophysiology of PTL. *G. vaginalis* and other anaerobes such as *Prevotella* spp. are capable of degrading cervical mucus, promoting ascending intrauterine infections, local inflammation, uterine motility, and PTL [33,34]. The treatment of bacterial vaginosis for all symptomatic pregnant women is recommended in many international guidelines, so as to reduce the risk of adverse pregnancy outcomes including PPROM and PTB [35,36]. Although controversial, some national guidelines also recommend screening for bacterial vaginosis and treatment of women with a history of PTB [36–39]. Our study identified *Gardnerella* spp. as risk factors for PTL; however, it is the absence of lactobacilli that appears to increase the risk of sPTB. These findings align with numerous metagenomic studies highlighting the protective role of lactobacilli, particularly *L. crispatus*, *L. gasseri*, and *L. jensenii* [19,22,40]. They support a comprehensive management of bacterial vaginosis in pregnancy aimed at restoring a healthy microbiota. Such a management strategy would likely combine antibiotics targeting anaerobes with probiotics, as has been proposed for the treatment of recurrent vaginosis [38].

Our results also raise questions about the management of vaginal colonization by enterobacteria during PTL and PPROM. In contrast to GBS, there are currently no consensus guidelines for the screening and administration of antibiotics against enterobacteria during pregnancy, even in the case of PPROM. In the absence of evidence of infection, treatment strategies vary widely between countries, ranging from no treatment to broad-spectrum antibiotics such as 3^rd^ generation cephalosporins [41]. Altogether, our observations underscore the necessity for studies to assess the efficacy of prophylaxis tailored to the pathogenic species present in the vaginal microbiota of women at high risk of preterm delivery.

The alignment of our findings with those of prior metagenomic studies demonstrates the reliability and practicality of vaginal microbiota monitoring through microbiological culture methods. Until metagenomics becomes widely adopted in routine clinical practice, routine conventional microbiological techniques could be used for risk stratification in women at risk of preterm delivery and for their follow-up. Microbiological culture results could be integrated into a decision-making algorithm alongside other clinical risk factors, such as gestational diabetes, or BMI. These techniques are indeed easily transposable and reproducible among clinical laboratories, possibly in combination with nucleic acid amplification tests (such as PCR tests) targeting specific pathogenic species. These tests are readily integrated into laboratory procedures and are already available for the diagnosis of bacterial vaginosis or the detection of GBS [42,43]. They could be adapted for the management of women at high risk for preterm delivery by means of additional detection of enterobacteria.

### Research Implications

Interventional studies to assess the efficacy and tolerability of antibiotics alone or in combination with probiotics to reduce the risk of PTB and PPROM in high-risk populations are still lacking. Randomized control trials should consider that a particular probiotic strain may not be protective for all women, according to their CST. The benefit of personalized targeted antibiotic prophylaxis directed against specific pathogenic species, e.g., GBS, enterobacteria, and *Candida* spp., in reducing the risk of PTB, especially in the case of PPROM, should also be investigated, together with their impact on maternal vaginal and intestinal microbiota, on the establishment of the neonatal microbiome, and on their long-term impact on infant health. There is also an important need to investigate the association between the vaginal microbiota and the risk of early-onset neonatal sepsis [44,45].

In addition, further studies are needed to decipher the role of the microbiota on pregnancy outcomes. Particularly, longitudinal studies aimed to investigate whether the observed changes in microbiota composition are a cause or a consequence of PTL and PPROM, together with mechanistic studies aimed to identify the underlying host-microbe molecular mechanisms are of utmost importance for the design of specific targeted interventions.

### Strengths and Limitations

This is a large cohort of over 1,800 patients, including over 400 PTL and nearly 400 PPROM along with low-risk controls. The sample size allowed adjustment for important confounders of the outcome measures which are often overlooked, including demographic factors, such as country of birth, BMI, history of PTB, and gestational diabetes. Hence, we were able to demonstrate that the vaginal microbiota composition represents an independent risk factor for PTL and PPROM, as well as for sPTB. In contrast to metagenomics, the conventional culture approach is widely available in microbiology laboratories and routine clinical settings, and thus easily reproducible and applicable to large-scale studies at lower costs.

However, there are limitations that need to be considered. First of all, we did not perform a longitudinal microbiota profiling across pregnancy, which would have added insights into temporal microbiome dynamics and their potential predictive value, especially for PTL and PPROM. Indeed, our main objective was to investigate the impact of the vaginal microbiota in women at high risk of sPTB, in a study population of sufficient sample size to allow the analysis of confounders. To overcome sample size limitations, we decided to enroll women presenting with PTL or PPROM rather than healthy pregnancies in their first trimester. Because of the study design, the vaginal samples were not all collected at the same time of gestation, mostly after 35 weeks of gestation for controls and before 35 weeks of gestation for women in the PTL and PPROM groups, at a median of 29 and 32 weeks, respectively. Yet, the microbiome is dynamic and changes during pregnancy [46]. Therefore, the differences in vaginal microbiota composition between the control group and the other two cohort groups may be due, at least in part, to differences in the timing of sampling. However, a previous study demonstrated that the abundance of lactobacilli is greater in the microbiomes of pregnant women with *Lactobacillus*-dominated CSTs in comparison with non-pregnant women as early as 11-16 weeks of gestation [47]. This suggests that the modifications of the vaginal microbiota, including increased lactobacilli abundance, appear early in pregnancy. In addition, other studies identified anaerobe-dominant and lactobacilli-dominant vaginal microbiota as early as 8-20 weeks of gestation, demonstrating that the formers were at increased risk of PTB [20,48]. Moreover, longitudinal studies demonstrate that vaginal microbiota composition remains stable in non-African women when comparing early and late pregnancy, i.e. before and after 32 weeks of gestation, respectively. However, in African women, there is a transition from taxa associated with dysbiosis to lactobacilli-dominated microbiota, specifically from CST-IV (non-*Lactobacillus*-dominated) to CST-I and CST-III (*L. crispatus*– and *Lactobacillus iners*-dominated microbiota, respectively) [12,15,49]. Nevertheless, these shifts occur predominantly before 24 weeks of gestation. Therefore, the observed differences between the two at-risk groups and the control group in our study are more likely to reflect actual microbial signatures of PTL and PPROM than differences related solely to the timing of sampling. Nevertheless, a longitudinal sampling investigating microbiota trajectory before and after enrollment, especially to determine its impact on pregnancy outcome, would have been particularly interesting.

A second limitation is that the microbiota analyses were performed using vaginal swab cultures. This leads to three main issues: *(i)* fastidious and unculturable species such as *Atopobium* spp., or Mollicutes could not be studied, thereby leading to the underestimation of microbiota diversity compared to sequencing methods; *(ii)* culture is less sensitive than metagenomics, and the reported species were most likely the dominant ones whereas minor species were likely overlooked; *(iii)* quantitative analysis could not be performed and only the presence or absence of major species was studied in statistical analyses. However, our results are consistent with those of metagenomic-based studies, particularly regarding the significance of *Gardnerella* spp. and lactobacilli in pregnancy outcomes [8]. Another limitation of the study is the lack of identification of lactobacilli at the species level. As a result, the vaginal microbiotas were not categorized according to CSTs, and the protective role of the different *Lactobacillus* species could not be addressed. Indeed, not all *Lactobacillus* species offer equal protection against PTB. *L. iners*, in particular, is increasingly regarded as a “false friend” due to the higher instability of the microbiomes dominated by this species compared to those dominated by *L. crispatus* [50,51]. Finally, although our study is multicenter and includes women from diverse origins, it is a French national cohort, and the results may be biased toward the White Caucasian ethnicity.

With regard to the analysis of confounders, residual confounders such as antibiotic use, sexual behavior, socioeconomic status, and prior microbiota-altering interventions could not be investigated, due to the diversity or complexity of the data, or because the data were missing. Furthermore, the microbiota culture status was defined based solely on microbiological culture results, and its interpretation requires caution. The clinical criteria for lower genital tract disorders, such as redness, burning, and vaginal discharge, for the diagnosis of aerobic vaginitis, the Amsel criteria for the clinical diagnosis of bacterial vaginosis, and vaginal pro-inflammatory cytokines were not included in the study, despite their demonstrated relevance in assessing pregnancy outcomes [52–55]. Indeed, the inclusion of an excessive number of variables in statistical models has the potential to result in errors and erroneous conclusions. For this reason, we restricted our analyses to the most robust variables.

## Conclusion

In this large cohort study, we used routine microbiological cultures and multivariable analyses, adjusting for confounding variables, to identify vaginal microbiota signatures of PTL and PPROM and risk factors for sPTB in these high-risk populations. Our findings confirm the protective role of lactobacilli during pregnancy and the association between *Gardnerella* spp. and PTL. Furthermore, we highlight enterobacteria as a risk factor for adverse outcomes and we demonstrate that, among women presenting with PTL or PPROM, a vaginal microbiota depleted of lactobacilli constitutes a strong risk factor for sPTB. While further studies are needed to determine whether interventions aimed at restoring a healthy microbiota could reduce adverse pregnancy outcomes, our results open new perspectives for conventional microbiological methods, including culture and molecular tests, for the management of women at high risk of PTL, PPROM, and PTB in routine clinical settings.

## Transparency declaration

### Ethics statement

The reported study was done in accordance with the declaration of Helsinki and reviewed by the French ethics Committee for Protection of Persons who granted approval on November, 4^th^ 2017 (identifier 2017 – A02755-48; NCT03371056). All individuals enrolled in the cohort provided signed informed consent that included authorization for the collection of medical information and biological samples. All collected data were anonymized and integrated in a secured electronic database.

## Data availability statement

All authors had full access to data. Data will be made available upon requests directed to the corresponding author.

## Acknowledgements

We thank all the contributors to this study, collectively called the InSPIRe Consortium. The full list can be found in the accompanying Supplementary Information.

Part of the results were previously presented in April 2023 at the ECCMID scientific meeting (Copenhagen, Denmark), in September 2023 at the ESPBC congress (Haarlem, The Netherlands), and in January 2025 at the SMFM 2025 pregnancy meeting (Denver, Colorado, USA).

## Authors’ contributions

Conceptualization: C.Pa., C.Po., L.M., and A.T. Methodology: J.R., C.Pl., L.La., F.G., P.-Y.A., C.Pa., L.M., and A.T. Formal analysis: L.Le., J.R., C.Pl., L.La., N.G., F.G., P.-Y.A., L.M., and A.T. Investigation: L.Le., J.R., and A.T. Resources: C.Pl., L.La, N.G., F.G., and L.M. Data curation: L.Le, J.R., C.Pl, L.La, N.G., and A.T. Writing – Original draft: L.Le and A.T. Writing – Review and Editing: all authors. Supervision: C.Pa, C.Po., L.M. and A.T. Funding acquisition: C.Pa, C.Po and L.M.

## Abbreviations

aOR: adjusted odds ratio
BMI: Body mass index
CI: Confidence interval
CST: Community state type
GBS: Group B *Streptococcus*
OR: Odds ratio
PPROM: Preterm premature rupture of membranes
PTB: Preterm birth
PTL: Preterm labor
sPTB: Spontaneous preterm birth

## Supplementary Material

**Supplementary Table 1.** Vaginal microbiota culture status according to the microbiological culture results.

**Supplementary Table 2.** Full list of microbial species identified and reported from vaginal swab cultures.

**Supplementary Table 3.** Correlation between the Nugent score and the microbiology culture status.

**Supplementary Table 4.** Nugent scores at enrollment according to the place of birth.

**Supplementary Table 5.** Microbiological cultures of vaginal samples at enrollment according to the place of birth.

**Supplementary Table 6.** Nugent score at enrollment according to gravidity and parity.

**Supplementary Table 7.** Microbiological cultures of vaginal samples at enrollment according to gravidity and parity.

**Supplementary Table 8.** Nugent scores at enrollment according to body mass index.

**Supplementary Table 9.** Microbiological cultures of vaginal samples at enrollment according to the body mass index.

**Supplementary Figure 1. Second model of multivariable analysis of risk factors for preterm labor and preterm premature rupture of membranes**.

In this model, *Escherichia coli* and *Klebsiella pneumoniae* were used instead of enterobacteria as variables. Error bars indicate the upper and lower limit of the 95% confidence interval (CI); aOR: adjusted odds ratio.^a^ aOR (95% CI) of Center 3 for preterm labor and preterm premature rupture of membranes: 1.6×10^8^ (6.7×10^2^-2.5×10^70^) and 2.4×10^8^ (0.09-6.3×10^85^), respectively.

**Supplementary Figure 2. Second model of multivariable analysis of risk factors for spontaneous preterm birth**.

In this model, *Escherichia coli* and *Klebsiella pneumoniae* were used instead of enterobacteria as variables. Error bars indicate the upper and lower limit of the 95% confidence interval (CI); aOR: adjusted odds ratio. WG: weeks of gestation.

**Supplementary Information. Composition of the Inspire consortium.**

## References

[1] E.O. Ohuma, A.-B. Moller, E. Bradley, S. Chakwera, L. Hussain-Alkhateeb, A. Lewin, Y.B. Okwaraji, W.R. Mahanani, E.W. Johansson, T. Lavin, D.E. Fernandez, G.G. Domínguez, A. de Costa, J.A. Cresswell, J. Krasevec, J.E. Lawn, H. Blencowe, J. Requejo, A.C. Moran, National, regional, and global estimates of preterm birth in 2020, with trends from 2010: a systematic analysis, Lancet 402 (2023) 1261–1271. 10.1016/S0140-6736(23)00878-4.

[2] M.S. Harrison, R.L. Goldenberg, Global burden of prematurity, Semin Fetal Neonatal Med 21 (2016) 74–79. 10.1016/j.siny.2015.12.007.

[3] GBD 2016 Causes of Death Collaborators, Global, regional, and national age-sex specific mortality for 264 causes of death, 1980-2016: a systematic analysis for the Global Burden of Disease Study 2016, Lancet 390 (2017) 1151–1210. 10.1016/S0140-6736(17)32152-9.

[4] R.L. Goldenberg, J.F. Culhane, J.D. Iams, R. Romero, Epidemiology and causes of preterm birth, Lancet 371 (2008) 75–84. 10.1016/S0140-6736(08)60074-4.

[5] E. Lorthe, Épidémiologie, facteurs de risque et pronostic de l’enfant. RPC: rupture prématurée des membranes avant terme CNGOF, Gynécologie Obstétrique Fertilité & Sénologie 46 (2018) 1004–1021. 10.1016/j.gofs.2018.10.019.

[6] E. Bayar, P.R. Bennett, D. Chan, L. Sykes, D.A. MacIntyre, The pregnancy microbiome and preterm birth, Semin Immunopathol 42 (2020) 487–499. 10.1007/s00281-020-00817-w.

[7] H.J. Schuster, A.M. Bos, L. Himschoot, R. Van Eekelen, S.P.F. Matamoros, M.A. De Boer, M.A. Oudijk, C. Ris-Stalpers, P. Cools, P.H.M. Savelkoul, R.C. Painter, R. Van Houdt, Vaginal microbiota and spontaneous preterm birth in pregnant women at high risk of recurrence, Heliyon 10 (2024) e30685. 10.1016/j.heliyon.2024.e30685.

[8] A.M. Powell, F.Z.A. Khan, J. Ravel, M.A. Elovitz, Untangling Associations of Microbiomes of Pregnancy and Preterm Birth, Clin Perinatol 51 (2024) 425–439. 10.1016/j.clp.2024.02.009.

[9] S. Greenbaum, G. Greenbaum, J. Moran-Gilad, A.Y. Weintraub, Ecological dynamics of the vaginal microbiome in relation to health and disease, Am J Obstet Gynecol 220 (2019) 324–335. 10.1016/j.ajog.2018.11.1089.

[10] S.D. Song, K.D. Acharya, J.E. Zhu, C.M. Deveney, M.R.S. Walther-Antonio, M.J. Tetel, N. Chia, Daily Vaginal Microbiota Fluctuations Associated with Natural Hormonal Cycle, Contraceptives, Diet, and Exercise, mSphere 5 (2020) e00593–20. 10.1128/mSphere.00593-20.

[11] J. Ravel, P. Gajer, Z. Abdo, G.M. Schneider, S.S.K. Koenig, S.L. McCulle, S. Karlebach, R. Gorle, J. Russell, C.O. Tacket, R.M. Brotman, C.C. Davis, K. Ault, L. Peralta, L.J. Forney, Vaginal microbiome of reproductive-age women, Proc. Natl. Acad. Sci. U.S.A. 108 (2011) 4680–4687. https://doi.org/bayar.

[12] R. Romero, K.R. Theis, N. Gomez-Lopez, A.D. Winters, J.J. Panzer, H. Lin, J. Galaz, J.M. Greenberg, Z. Shaffer, D.J. Kracht, T. Chaiworapongsa, E. Jung, F. Gotsch, J. Ravel, S.D. Peddada, A.L. Tarca, The Vaginal Microbiota of Pregnant Women Varies with Gestational Age, Maternal Age, and Parity, Microbiol Spectr 11 (2023) e03429–22. 10.1128/spectrum.03429-22.

[13] K. Kervinen, T. Holster, S. Saqib, S. Virtanen, V. Stefanovic, L. Rahkonen, P. Nieminen, A. Salonen, I. Kalliala, Parity and gestational age are associated with vaginal microbiota composition in term and late term pregnancies, eBioMedicine 81 (2022) 104107. 10.1016/j.ebiom.2022.104107.

[14] M.T. France, B. Ma, P. Gajer, S. Brown, M.S. Humphrys, J.B. Holm, L.E. Waetjen, R.M. Brotman, J. Ravel, VALENCIA: a nearest centroid classification method for vaginal microbial communities based on composition, Microbiome 8 (2020) 166. 10.1186/s40168-020-00934-6.

[15] R. Romero, S.S. Hassan, P. Gajer, A.L. Tarca, D.W. Fadrosh, L. Nikita, M. Galuppi, R.F. Lamont, P. Chaemsaithong, J. Miranda, T. Chaiworapongsa, J. Ravel, The composition and stability of the vaginal microbiota of normal pregnant women is different from that of non-pregnant women, Microbiome 2 (2014) 4. 10.1186/2049-2618-2-4.

[16] D.A. MacIntyre, M. Chandiramani, Y.S. Lee, L. Kindinger, A. Smith, N. Angelopoulos, B. Lehne, S. Arulkumaran, R. Brown, T.G. Teoh, E. Holmes, J.K. Nicoholson, J.R. Marchesi, P.R. Bennett, The vaginal microbiome during pregnancy and the postpartum period in a European population, Sci Rep 5 (2015) 8988. 10.1038/srep08988.

[17] J.M. Fettweis, M.G. Serrano, J.P. Brooks, D.J. Edwards, P.H. Girerd, H.I. Parikh, B. Huang, T.J. Arodz, L. Edupuganti, A.L. Glascock, J. Xu, N.R. Jimenez, S.C. Vivadelli, S.S. Fong, N.U. Sheth, S. Jean, V. Lee, Y.A. Bokhari, A.M. Lara, S.D. Mistry, R.A. Duckworth, S.P. Bradley, V.N. Koparde, X.V. Orenda, S.H. Milton, S.K. Rozycki, A.V. Matveyev, M.L. Wright, S.V. Huzurbazar, E.M. Jackson, E. Smirnova, J. Korlach, Y.-C. Tsai, M.R. Dickinson, J.L. Brooks, J.I. Drake, D.O. Chaffin, A.L. Sexton, M.G. Gravett, C.E. Rubens, N.R. Wijesooriya, K.D. Hendricks-Muñoz, K.K. Jefferson, J.F. Strauss, G.A. Buck, The vaginal microbiome and preterm birth, Nat Med 25 (2019) 1012–1021. 10.1038/s41591-019-0450-2.

[18] U. Gudnadottir, J.W. Debelius, J. Du, L.W. Hugerth, H. Danielsson, I. Schuppe-Koistinen, E. Fransson, N. Brusselaers, The vaginal microbiome and the risk of preterm birth: a systematic review and network meta-analysis, Sci Rep 12 (2022) 7926. 10.1038/s41598-022-12007-9.

[19] A. Baud, K.-H. Hillion, C. Plainvert, V. Tessier, A. Tazi, L. Mandelbrot, C. Poyart, S.P. Kennedy, Microbial diversity in the vaginal microbiota and its link to pregnancy outcomes, Sci Rep 13 (2023) 9061. 10.1038/s41598-023-36126-z.

[20] M.A. Elovitz, P. Gajer, V. Riis, A.G. Brown, M.S. Humphrys, J.B. Holm, J. Ravel, Cervicovaginal microbiota and local immune response modulate the risk of spontaneous preterm delivery, Nat Commun 10 (2019) 1305. 10.1038/s41467-019-09285-9.

[21] K. Gorczyca, M.M. Kozioł, Ż. Kimber-Trojnar, J. Kępa, M. Satora, A.K. Rekowska, B. Leszczyńska-Gorzelak, Premature rupture of membranes and changes in the vaginal microbiome – Probiotics, Reproductive Biology 24 (2024) 100899. 10.1016/j.repbio.2024.100899.

[22] B.J. Callahan, D.B. DiGiulio, D.S.A. Goltsman, C.L. Sun, E.K. Costello, P. Jeganathan, J.R. Biggio, R.J. Wong, M.L. Druzin, G.M. Shaw, D.K. Stevenson, S.P. Holmes, D.A. Relman, Replication and refinement of a vaginal microbial signature of preterm birth in two racially distinct cohorts of US women, Proc. Natl. Acad. Sci. U.S.A. 114 (2017) 9966–9971. 10.1073/pnas.1705899114.

[23] J. Liao, L. Shenhav, J.A. Urban, M. Serrano, B. Zhu, G.A. Buck, T. Korem, Microdiversity of the vaginal microbiome is associated with preterm birth, Nat Commun 14 (2023) 4997. 10.1038/s41467-023-40719-7.

[24] HAS, Prévention anténatale du risque infectieux bactérien néonatal précoce, (2001). https://www.has-sante.fr/upload/docs/application/pdf/prevention_antenatale_du_risque_infectieux_bacterien_-_rec.pdf.

[25] H. Madar, Prise en charge thérapeutique (hors antibiothérapie) de la rupture prématurée des membranes avant terme. RPC rupture prématurée des membranes avant terme CNGOF. Management of preterm premature rupture of membranes (except for antibiotherapy): CNGOF preterm premature rupture of membranes guidelines., Gynécologie Obstétrique Fertilité & Sénologie 46 (2018) 1029–1042. 10.1016/j.gofs.2018.10.020.

[26] L.M. Filkins, O.B. Garner, Genital Cultures, in: ClinMicroNow, 1st ed., Wiley, 2023: pp. 1–46. 10.1002/9781683670438.cmph0011.

[27] M. De La Rosa, M. Perez, C. Carazo, L. Pareja, J.I. Peis, F. Hernandez, New Granada Medium for detection and identification of group B streptococci, J Clin Microbiol 30 (1992) 1019–1021. 10.1128/jcm.30.4.1019-1021.1992.

[28] L.F. Westblade, O.B. Garner, K. MacDonald, C. Bradford, D.H. Pincus, A.B. Mochon, R. Jennemann, R. Manji, M. Bythrow, M.A. Lewinski, C.-A.D. Burnham, C.C. Ginocchio, Assessment of Reproducibility of Matrix-Assisted Laser Desorption Ionization–Time of Flight Mass Spectrometry for Bacterial and Yeast Identification, J Clin Microbiol 53 (2015) 2349–2352. 10.1128/JCM.00187-15.

[29] R.P. Nugent, M.A. Krohn, S.L. Hillier, Reliability of diagnosing bacterial vaginosis is improved by a standardized method of gram stain interpretation, J Clin Microbiol 29 (1991) 297–301. 10.1128/jcm.29.2.297-301.1991.

[30] R.G. Brown, M. Al-Memar, J.R. Marchesi, Y.S. Lee, A. Smith, D. Chan, H. Lewis, L. Kindinger, V. Terzidou, T. Bourne, P.R. Bennett, D.A. MacIntyre, Establishment of vaginal microbiota composition in early pregnancy and its association with subsequent preterm prelabor rupture of the fetal membranes, Translat Res 207 (2019) 30–43. 10.1016/j.trsl.2018.12.005.

[31] S.L. Hillier, J. Martius, M. Krohn, N. Kiviat, K.K. Holmes, D.A. Eschenbach, A Case–Control Study of Chorioamnionic Infection and Histologic Chorioamnionitis in Prematurity, N Engl J Med 319 (1988) 972–978. 10.1056/NEJM198810133191503.

[32] D.A. Eschenbach, M.G. Gravett, K.C. Chen, U.B. Hoyme, K.K. Holmes, Bacterial vaginosis during pregnancy. An association with prematurity and postpartum complications, Scand J Urol Nephrol Suppl 86 (1984) 213–222.

[33] L. Howe, R. Wiggins, P.W. Soothill, M.R. Millar, P.J. Horner, A.P. Corfield, Mucinase and sialidase activity of the vaginal microflora: implications for the pathogenesis of preterm labour, Int J STD AIDS 10 (1999) 442–447. 10.1258/0956462991914438.

[34] W.G. Lewis, L.S. Robinson, N.M. Gilbert, J.C. Perry, A.L. Lewis, Degradation, Foraging, and Depletion of Mucus Sialoglycans by the Vagina-adapted Actinobacterium Gardnerella vaginalis, J Biol Chem 288 (2013) 12067–12079. 10.1074/jbc.M113.453654.

[35] K.A. Workowski, L.H. Bachmann, P.A. Chan, C.M. Johnston, C.A. Muzny, I. Park, H. Reno, J.M. Zenilman, G.A. Bolan, Sexually Transmitted Infections Treatment Guidelines, 2021, MMWR Recomm. Rep. 70 (2021) 1–187. 10.15585/mmwr.rr7004a1.

[36] Collège national des gynécologues et obstétriciens français, Prévention de la prématurité spontanée et de ses conséquences, (2016). https://cngof.fr/app/pdf/RPC//RPC%20DU%20CNGOF/2016/RPC_2016_Prmaturit_spontane.pdf?x29325.

[37] British Association for Sexual Health and HIV, Bacterial Vaginosis, (2012). https://www.bashh.org/resources/21/bacterial_vaginosis_2012/.

[38] A. Farr, S. Swidsinski, D. Surbek, B.F. Tirri, B. Willinger, U. Hoyme, G. Walter, I. Reckel-Botzem, W. Mendling, Bacterial Vaginosis: Guideline of the DGGG, OEGGG and SGGG (S2k-Level, AWMF Registry No. 015/028, June 2023), Geburtshilfe Frauenheilkd 83 (2023) 1331–1349. 10.1055/a-2169-8539.

[39] D. Subtil, G. Brabant, E. Tilloy, P. Devos, F. Canis, A. Fruchart, M.-C. Bissinger, J.-C. Dugimont, C. Nolf, C. Hacot, S. Gautier, J. Chantrel, M. Jousse, D. Desseauve, J.L. Plennevaux, C. Delaeter, S. Deghilage, A. Personne, E. Joyez, E. Guinard, E. Kipnis, K. Faure, B. Grandbastien, P.-Y. Ancel, F. Goffinet, R. Dessein, Early clindamycin for bacterial vaginosis in pregnancy (PREMEVA): a multicentre, double-blind, randomised controlled trial, Lancet 392 (2018) 2171–2179. 10.1016/S0140-6736(18)31617-9.

[40] N. Tabatabaei, A. Eren, L. Barreiro, V. Yotova, A. Dumaine, C. Allard, W. Fraser, Vaginal microbiome in early pregnancy and subsequent risk of spontaneous preterm birth: a case–control study, BJOG 126 (2019) 349–358. 10.1111/1471-0528.15299.

[41] D.E. Werter, I. Dehaene, L. Gurney, M. Vargas Buján, B.M. Kazemier, Differences in clinical practice regarding screening and treatment of infections associated with spontaneous preterm birth: An international survey, Eur J Obstet Gynecol Reprod Biol 266 (2021) 83–88. 10.1016/j.ejogrb.2021.09.009.

[42] R.F. Lamont, E.H. Van Den Munckhof, B.M. Luef, C.A. Vinter, J.S. Jørgensen, Recent advances in cultivation-independent molecular-based techniques for the characterization of vaginal eubiosis and dysbiosis, Fac Rev 9 (2020). 10.12703/r/9-21.

[43] J. Peng, Y. Liu, J. Zou, J. Wang, C.D. Jorge Luis, H. Zhong, Accuracy of real-time polymerase chain reaction test for Group B *Streptococcus* detection in pregnant women: A systematic review and meta-analysis, Eur J Obstet Gynecol Reprod Biol 304 (2025) 141–151. 10.1016/j.ejogrb.2024.11.035.

[44] I. Ben M’Barek, L. Landraud, L. Desfrere, K. Sallah, C. Couffignal, M. Schneider, L. Mandelbrot, Contribution of vaginal culture to predict early onset neonatal infection in preterm prelabor rupture of membranes, Eur J Obstet Gynecol Reprod Biol 261 (2021) 78–84. 10.1016/j.ejogrb.2021.04.016.

[45] L. Mandelbrot, S. Kennedy, J. Rousseau, F. Goffinet, L. Landraud, C. Plainvert, V. Marcou, L. Desfrère, T. Barral, L. Allal, A. Baud, N. Grall, C. Poyart, P.-Y. Ancel, A. Tazi, Predicting neonatal infection in preterm premature rupture of membranes with vaginal microbiology and metagenomics: a prospective cohort study, Am J Obstet Gynecol (2025) S0002937825009391. 10.1016/j.ajog.2025.12.042.

[46] S. Onyango, J.D. Mi, A. Koech, P. Okiro, M. Temmerman, P. Von Dadelszen, R.M. Tribe, G. Omuse, the PRECISE Network, Microbiota dynamics, metabolic and immune interactions in the cervicovaginal environment and their role in spontaneous preterm birth, Front. Immunol. 14 (2023) 1306473. 10.3389/fimmu.2023.1306473.

[47] A.C. Freitas, B. Chaban, A. Bocking, M. Rocco, S. Yang, J.E. Hill, D.M. Money, The VOGUE Research Group, S. Hemmingsen, G. Reid, T. Dumonceaux, G. Gloor, M. Links, K. O’Doherty, P. Tang, J. Van Schalkwyk, M. Yudin, The vaginal microbiome of pregnant women is less rich and diverse, with lower prevalence of Mollicutes, compared to non-pregnant women, Sci Rep 7 (2017) 9212. 10.1038/s41598-017-07790-9.

[48] A.L. Dunlop, G.A. Satten, Y.-J. Hu, A.K. Knight, C.C. Hill, M.L. Wright, A.K. Smith, T.D. Read, B.D. Pearce, E.J. Corwin, Vaginal Microbiome Composition in Early Pregnancy and Risk of Spontaneous Preterm and Early Term Birth Among African American Women, Front. Cell. Infect. Microbiol. 11 (2021) 641005. 10.3389/fcimb.2021.641005.

[49] M.G. Serrano, H.I. Parikh, J.P. Brooks, D.J. Edwards, T.J. Arodz, L. Edupuganti, B. Huang, P.H. Girerd, Y.A. Bokhari, S.P. Bradley, J.L. Brooks, M.R. Dickinson, J.I. Drake, R.A. Duckworth, S.S. Fong, A.L. Glascock, S. Jean, N.R. Jimenez, J. Khoury, V.N. Koparde, A.M. Lara, V. Lee, A.V. Matveyev, S.H. Milton, S.D. Mistry, S.K. Rozycki, N.U. Sheth, E. Smirnova, S.C. Vivadelli, N.R. Wijesooriya, J. Xu, P. Xu, D.O. Chaffin, A.L. Sexton, M.G. Gravett, C.E. Rubens, K.D. Hendricks-Muñoz, K.K. Jefferson, J.F. Strauss, J.M. Fettweis, G.A. Buck, Racioethnic diversity in the dynamics of the vaginal microbiome during pregnancy, Nat Med 25 (2019) 1001–1011. 10.1038/s41591-019-0465-8.

[50] L.M. Kindinger, P.R. Bennett, Y.S. Lee, J.R. Marchesi, A. Smith, S. Cacciatore, E. Holmes, J.K. Nicholson, T.G. Teoh, D.A. MacIntyre, The interaction between vaginal microbiota, cervical length, and vaginal progesterone treatment for preterm birth risk, Microbiome 5 (2017) 6. 10.1186/s40168-016-0223-9.

[51] H. Verstraelen, R. Verhelst, G. Claeys, E. De Backer, M. Temmerman, M. Vaneechoutte, Longitudinal analysis of the vaginal microflora in pregnancy suggests that *L. crispatus* promotes the stability of the normal vaginal microflora and that *L. gasseri* and/or *L. iners* are more conducive to the occurrence of abnormal vaginal microflora, BMC Microbiol 9 (2009) 116. 10.1186/1471-2180-9-116.

[52] S. Hadhoum, D. Subtil, J. Labreuche, E. Couvreur, G. Brabant, R. Dessein, R. Le Guern, Reassessing the association between bacterial vaginosis and preterm birth: A systematic review and meta-analysis, J Gynecol Obstet Hum Reprod 54 (2025) 102871. 10.1016/j.jogoh.2024.102871.

[53] G. Donders, K. Van Calsteren, G. Bellen, R. Reybrouck, T. Van Den Bosch, I. Riphagen, S. Van Lierde, Predictive value for preterm birth of abnormal vaginal flora, bacterial vaginosis and aerobic vaginitis during the first trimester of pregnancy, BJOG 116 (2009) 1315–1324. 10.1111/j.1471-0528.2009.02237.x.

[54] R. Romero, S.K. Dey, S.J. Fisher, Preterm labor: One syndrome, many causes, Science 345 (2014) 760–765. 10.1126/science.1251816.

[55] G. Donders, G. Bellen, S. Grinceviciene, K. Ruban, P. Vieira-Baptista, Aerobic vaginitis: no longer a stranger, Res Microbiol 168 (2017) 845–858. 10.1016/j.resmic.2017.04.004.

