## Supplemental Information for "Culture-based analysis of vaginal microbiota and its link to pregnancy outcomes: an observational cohort study in France"

### InSPIRE Consortium


Microbiology

**Clermont Olivier** Inserm 1137 IAME Université Paris Cité

**Nathalie Grall** APHP Bichat Microbiology

Methodology

**Pierre-Yves Ancel**, APHP URC-CIC Cochin-Necker Mère-Enfant – U1153 EPOPé, FHU Prema

**Laurence Lecomte**, APHP DRCI, FHU Prema

**Hendy Abdoul**, APHP URC-CIC Cochin-Necker Mère-Enfant, FHU Prema

**Jessica Rousseau**, Data Manager, APHP URC-CIC Cochin-Necker Mère-Enfant

Obstetrics

**Laurent Mandelbrot**, APHP Louis Mourier Obstetrics/Gynecology, FHU Prema, Inserm 1137 IAME Université Paris Cité

**François Goffinet**, APHP Cochin Port-Royal Obstetrics/Gynecology, FHU Prema, U1153 EPOPé, Université Paris Cité

**Dominique Luton** APHP Bichat Obstetrics/Gynecology, Université Paris Cité

Neonatalogy

**Pierre-Henri Jarreau**, APHP Cochin Port-Royal, Neonatalogy, FHU Prema, Université Paris Cité

**Luc Desfrère**, APHP Louis Mourier Neonatalogy, FHU Prema

**Lahçene Allal**, APHP Bichat Neonatalogy, FHU Prema

Metagenomics

**Sean Kennedy**, Institut Pasteur, Université Paris Cité, Département de biologie computationnelle, F-75015 Paris, France

**Agnes Baud**, Institut Past, Institut Pasteur, Université Paris Cité, Département de biologie computationnelle, F-75015 Paris, France

**Kenzo-Hugo Hillion**, Institut Past, Institut Pasteur, Université Paris Cité, Département de biologie computationnelle, F-75015 Paris, France

Genomics

**Céline Méhats**, Institut Cochin Inserm U 1016, UMR CNRS 8104, Université Paris Cité, « Des gamètes à la naissance », Génomique, épigénétique et physiopathologie de la reproduction, FHU Prema

**Frédéric Batteux** Institut Cochin Inserm U 1016, UMR CNRS 8104, Université Paris Cité, « Des gamètes à la naissance », - Stress oxydant, prolifération cellulaire et inflammation, FHU Prema

Program manager

**Véronique Tessier**, APHP DRCI

Data monitoring and management

**Sinthiya Sivanesan**, APHP URC/CIC Necker Cochin, FHU Prema

**Hélène Jabbarian**, APHP Louis Mourier, URC Paris-Nord, FHU Prema

Industrial partner

**Christophe Pannetier**, BforCure

**Laura Lesimple**, Université Paris Cité, Inserm U1016 CNRS UMR 8104 Institut Cochin team Bacteria and Perinatality, BforCure
