## Supplemental Tables for "Culture-based analysis of vaginal microbiota and its link to pregnancy outcomes: an observational cohort study in France"

**Supplementary Table 1.** Vaginal microbiota culture status according to the microbiological culture results.

| **Lactobacilli** | **Aerobic bacteria**[a] | **Anaerobic bacteria**[b] | **Candida spp.** | **Microbiota culture status** |
|---|---|---|---|---|
| Numerous | Absent - Rare | Absent - Rare | Absent - Rare | Normal |
| Rare | Absent | Absent | Absent - Rare | Normal |
| Numerous | Numerous | Absent - Rare | Absent - Rare | Intermediate |
| Numerous | Absent - Rare | Numerous | Absent - Rare | Intermediate |
| Numerous | Numerous | Numerous | Absent - Rare | Intermediate |
| Rare | Rare | Absent | Absent - Rare | Intermediate |
| Rare | Absent | Rare | Absent - Rare | Intermediate |
| Rare | Rare | Rare | Absent - Rare | Intermediate |
| Absent - Rare | Numerous | Absent - Rare | Absent - Rare | Aerobic vaginitis / Mixed vaginitis[c] |
| Absent - Rare | Absent - Rare | Numerous | Absent - Rare | Anaerobes-dominated |
| Absent - Rare | Numerous | Numerous | Absent - Rare | Anaerobes-dominated |
| Absent - Rare - Numerous | Absent - Rare - Numerous | Absent - Rare - Numerous | Numerous | Candida |

[a] Including *Streptococcus* spp.
[b] Including *Gardnerella* spp.
[c] Aerobic vaginitis if dominated by a single bacterial species, mixed vaginitis if dominated by two or more aerobic bacterial species.

**Supplementary Table 2.** Full list of microbial species identified and reported from vaginal swab cultures.

| | Positive vaginal samples (N=1848) | |
|---|---|---|
| **Microbial species** | **Number** | **Percentage** |
| **Gram positive cocci** | | |
| *Enterococcus faecalis* | 620 | 33,55 % |
| *Enterococcus faecium* | 3 | 0,16 % |
| *Enterococcus hirae* | 1 | 0,05 % |
| *Micrococcus* sp. | 2 | 0,11 % |
| *Pediococcus acidilactici* | 1 | 0,05 % |
| *Staphylococcus aureus* | 93 | 5,03 % |
| *Staphylococcus condimenti* | 4 | 0,22 % |
| *Staphylococcus epidermidis* | 32 | 1,73 % |
| *Staphylococcus haemolyticus* | 25 | 1,35 % |
| *Staphylococcus hominis* | 6 | 0,32 % |
| *Staphylococcus lugdunensis* | 10 | 0,54 % |
| *Staphylococcus simulans* | 1 | 0,05 % |
| *Staphylococcus* sp. | 1 | 0,05 % |
| *Staphylococcus warneri* | 1 | 0,05 % |
| *Streptococcus agalactiae* (Group B *Streptococcus*) | 275 | 14,88 % |
| *Streptococcus anginosus* | 77 | 4,17 % |
| *Streptococcus dysgalactiae* | 1 | 0,05 % |
| *Streptococcus gallolyticus* | 4 | 0,22 % |
| *Streptococcus mitis* | 2 | 0,11 % |
| *Streptococcus oralis* | 3 | 0,16 % |
| *Streptococcus parasanguinis* | 1 | 0,05 % |
| *Streptococcus pasteurianus* | 3 | 0,16 % |
| *Streptococcus porcinus* | 2 | 0,11 % |
| *Streptococcus pseudoporcinus* | 1 | 0,05 % |
| *Streptococcus pyogenes* (Group A *Streptococcus*) | 1 | 0,05 % |
| *Streptococcus urinalis* | 3 | 0,16 % |
| *Streptococcus vestibularis* | 1 | 0,05 % |
| **Gram positive bacilli** | | |
| *Bacillus cereus* | 3 | 0,16 % |
| *Bacillus megaterium* | 4 | 0,22 % |
| *Bacillus simplex* | 1 | 0,05 % |
| *Bacillus* sp. | 8 | 0,43 % |
| *Brevibacterium ravenspurgense* | 1 | 0,05 % |
| *Corynebacterium aurimucosum* | 2 | 0,11 % |
| *Corynebacterium freneyi* | 1 | 0,05 % |
| *Corynebacterium striatum* | 2 | 0,11 % |
| *Corynebacterium* sp. | 9 | 0,49 % |
| *Lactobacillus* sp. | 1412 | 76,41 % |
| *Rothia mucilaginosa* | 1 | 0,05 % |
| **Gram negative bacilli** | | |
| Enterobacteria | | |
| *Citrobacter braakii* | 1 | 0,05 % |
| *Citrobacter freundii* | 2 | 0,11 % |
| *Citrobacter koseri* | 13 | 0,70 % |
| *Enterobacter asburiae* | 1 | 0,05 % |

| | Positive vaginal samples (N=1848) | |
|---|---|---|
| **Microbial species** | **Number** | **Percentage** |
| *Enterobacter cloacae* | 9 | 0,49 % |
| *Escherichia coli* | 304 | 16,45 % |
| *Hafnia alvei* | 1 | 0,05 % |
| *Klebsiella aerogenes* | 10 | 0,54 % |
| *Klebsiella oxytoca* | 5 | 0,27 % |
| *Klebsiella pneumoniae* | 82 | 4,44 % |
| *Mixta calida* | 1 | 0,05 % |
| *Morganella morganii* | 16 | 0,87 % |
| *Pantoea agglomerans* | 1 | 0,05 % |
| *Pantoea septica* | 2 | 0,11 % |
| *Pantoea* sp. | 1 | 0,05 % |
| *Proteus mirabilis* | 43 | 2,33 % |
| *Proteus vulgaris* | 3 | 0,16 % |
| *Raoultella* sp. | 1 | 0,05 % |
| *Serratia marcescens* | 2 | 0,11 % |
| Other Gram negative bacilli | | |
| *Acinetobacter baumannii* | 8 | 0,43 % |
| *Acinetobacter radioresistans* | 1 | 0,05 % |
| *Acinetobacter* sp. | 2 | 0,11 % |
| *Acinetobacter ursingii* | 2 | 0,11 % |
| *Haemophilus haemolyticus* | 1 | 0,05 % |
| *Haemophilus influenzae* | 19 | 1,03 % |
| *Haemophilus parainfluenzae* | 15 | 0,81 % |
| *Moraxella osloensis* | 4 | 0,22 % |
| *Moraxella* sp. | 1 | 0,05 % |
| *Pseudomonas aeruginosa* | 9 | 0,49 % |
| *Pseudomonas* sp. | 2 | 0,11 % |
| *Stenotrophomonas maltophilia* | 3 | 0,16 % |
| **Strict anaerobes** | | |
| *Actinomyces neuii* | 14 | 0,76 % |
| *Actinomyces* sp. | 1 | 0,05 % |
| *Actinomyces urogenitalis* | 1 | 0,05 % |
| *Alloscardovia omnicolens* | 14 | 0,76 % |
| *Alloscardovia* sp. | 2 | 0,11 % |
| *Bacteroides ovatus* | 1 | 0,05 % |
| *Bifidobacterium breve* | 13 | 0,70 % |
| *Bifidobacterium* sp. | 1 | 0,05 % |
| *Clostridium perfringens* | 1 | 0,05 % |
| *Finegoldia magna* | 6 | 0,32 % |
| *Gardnerella* spp. | 197 | 10,66 % |
| *Peptoniphilus harei* | 5 | 0,27 % |
| *Peptoniphilus indolicus* | 19 | 1,03 % |
| *Prevotella bivia* | 16 | 0,87 % |
| *Prevotella disiens* | 4 | 0,22 % |
| *Prevotella* sp. | 3 | 0,16 % |
| *Prevotella timonensis* | 1 | 0,05 % |
| *Veillonella atypica* | 6 | 0,32 % |
| *Veillonella* sp. | 1 | 0,05 % |
| *Veillonella rati* | 3 | 0,16 % |
| **Yeasts** | | |
| *Candida albicans* | 257 | 13,91 % |

| | Positive vaginal samples (N=1848) | |
|---|---|---|
| **Microbial species** | **Number** | **Percentage** |
| *Candida dubliniensis* | 1 | 0,05 % |
| *Candida glabrata* | 15 | 0,81 % |
| *Candida kefyr* | 3 | 0,16 % |
| *Candida krusei* | 7 | 0,38 % |
| *Candida lusitaniae* | 2 | 0,11 % |
| *Candida* sp. | 36 | 1,95 % |
| *Candida tropicalis* | 2 | 0,11 % |
| *Saccharomyces cerevisiae* | 1 | 0,05 % |

**Supplementary Table 3.** Correlation between the Nugent score and the microbiology culture status.

| **Nugent score** | **Normal (0-3)** | **Intermediate (4-6)** | **Bacterial vaginosis (≥7)** | **Total** |
|---|---|---|---|---|
| **Microbiology culture status, n (%)** | | | | |
| *Lactobacillus*-dominated | 925 (77.4) | 68 (36.8) | 6 (16.2) | 999 (70.5) |
| Aerobic vaginitis[a] | 58 (4.8) | 14 (7.6) | 3 (8.1) | 75 (5.3) |
| Mixed vaginitis[b] | 18 (1.5) | 6 (3.2) | 2 (5.4) | 26 (1.8) |
| Intermediate | 152 (12.7) | 67 (36.2) | 7 (18.9) | 226 (15.9) |
| Anaerobes-dominated[c] | 27 (2.3) | 27 (14.6) | 18 (48.7) | 72 (5.1) |
| *Candida*-dominated | 15 (1.3) | 3 (1.6) | 1 (2.7) | 19 (1.4) |
| *Missing* | *13 (1.1)* | *5 (2.6)* | *0 (0)* | *18 (1.3)* |
| **Total** | **1208 (84.2)** | **190 (13.2)** | **37 (2.6)** | **1435 (100)** |

[a] including *Streptococcus* spp.

[b] Aerobic vaginitis if dominated by a single bacterial species, mixed vaginitis if dominated by two or more aerobic bacterial species.

[c] including *Gardnerella* spp.

**Supplementary Table 4.** Nugent scores at enrollment according to the place of birth.

| **Nugent score, n (%)** | **France N=819** | **France, overseas territories[a] N=78** | **Europe, other countries N=121** | **North Africa N=389** | **Sub-Saharan Africa N=264** | **Asia N=64** | ***P* value[b]** |
|---|---|---|---|---|---|---|---|
| Normal (0-3) | 600 (86.3) | 34 (63.0) | 81 (81.8) | 227 (87.0) | 141 (78.3) | 45 (90.0) | <0.0001 |
| Intermediate (4-6) | 87 (12.5) | 9 (16.7) | 15 (15.2) | 30 (11.5) | 30 (16.7) | 4 (8.0) | |
| Bacterial vaginosis (≥7) | 8 (1.2) | 11 (20.4) | 3 (3.0) | 4 (1.5) | 9 (5.0) | 1 (2.0) | |
| *Missing* | *124 (15.1)* | *24 (30.8)* | *22 (18.2)* | *128 (32.9)* | *84 (31.8)* | *14 (21.9)* | |

[a] Including territories in South America (French Guiana), the Atlantic Ocean (Guadeloupe, Martinique, Saint-Martin, Saint-Barthélemy, Saint-Pierre-et-Miquelon), the Indian Ocean (Reunion Island, Mayotte), the Pacific Ocean (New Caledonia, French Polynesia, the Wallis and Futuna Islands), and the French Southern and Antarctic Territories.
[b] Chi-square test.

**Supplementary Table 5.** Microbiological cultures of vaginal samples at enrollment according to the place of birth.

| **Microbiological taxa, n (%)** | **France N=819** | **France, overseas territories[a] N=78** | **Europe, other countries N=121** | **North Africa N=389** | **Sub-Saharan Africa N=264** | **Asia N=64** | ***P* value[c]** |
|---|---|---|---|---|---|---|---|
| ***Lactobacillus* spp.** | | | | | | | <0.0001 |
| Numerous | 660 (81.4) | 41 (53.2) | 91 (75.2) | 252 (65.3) | 150 (57.9) | 51 (81.0) | |
| Rare | 28 (3.5) | 5 (6.5) | 6 (5.0) | 23 (6.0) | 14 (5.4) | 3 (4.8) | |
| Absent | 123 (15.2) | 31 (40.3) | 24 (19.8) | 111 (28.8) | 95 (36.7) | 9 (14.3) | |
| *Missing* | *8 (1.0)* | *1 (1.3)* | *0 (0)* | *3 (0.8)* | *5 (1.9)* | *1 (1.6)* | |
| **Enterobacteria, all species** | | | | | | | <0.0001 |
| Numerous | 92 (11.2) | 7 (9.0) | 11 (9.1) | 57 (14.7) | 48 (18.2) | 9 (14.1) | |
| Rare | 56 (6.8) | 11 (14.1) | 24 (19.8) | 40 (10.3) | 23 (8.7) | 9 (14.1) | |
| Absent | 670 (81.9) | 60 (76.9) | 86 (71.1) | 292 (75.1) | 193 (73.1) | 46 (71.9) | |
| *Missing* | *1 (0.1)* | *0 (0)* | *0 (0)* | *0 (0)* | *0 (0)* | *0 (0)* | |
| ***Escherichia coli*** | | | | | | | 0.0058 |
| Numerous | 70 (8.6) | 4 (5.1) | 6 (5.0) | 39 (10.1) | 27 (10.2) | 7 (10.9) | |
| Rare | 52 (6.4) | 6 (7.7) | 20 (16.5) | 25 (6.4) | 16 (6.1) | 9 (14.1) | |
| Absent | 694 (85.0) | 68 (87.2) | 95 (78.5) | 324 (83.5) | 221 (83.7) | 48 (75.0) | |
| *Missing* | *3 (0.4)* | *0 (0)* | *0 (0)* | *1 (0.3)* | *0 (0)* | *0 (0)* | |
| ***Klebsiella pneumoniae*** | | | | | | | <0.0001 |
| Numerous | 11 (1.3) | 5 (6.4) | 2 (1.7) | 16 (4.1) | 18 (6.8) | 0 (0) | |
| Rare | 4 (0.5) | 2 (2.6) | 2 (1.7) | 10 (2.6) | 6 (2.3) | 0 (0) | |
| Absent | 801 (98.2) | 71 (91.0) | 117 (96.7) | 362 (93.3) | 240 (90.9) | 64 (100) | |
| *Missing* | *3 (0.4)* | *0 (0)* | *0 (0)* | *1 (0.3)* | *0 (0)* | *0 (0)* | |
| ***Staphylococcus aureus*** | | | | | | | <0.0001 |
| Numerous | 15 (1.9) | 3 (3.8) | 1 (0.8) | 10 (2.6) | 17 (6.5) | 1 (1.6) | |
| Rare | 15 (1.9) | 0 (0) | 3 (2.5) | 6 (1.5) | 14 (5.3) | 2 (3.2) | |
| Absent | 774 (96.3) | 75 (96.2) | 117 (96.7) | 372 (95.9) | 232 (88.2) | 60 (95.2) | |
| *Missing* | *15 (1.8)* | *0 (0)* | *0 (0)* | *1 (0.3)* | *1 (0.4)* | *1 (1.6)* | |
| **Group B *Streptococcus*** | | | | | | | 0.061 |
| Numerous | 91 (11.1) | 6 (7.7) | 13 (10.7) | 51 (13.1) | 41 (15.5) | 9 (14.1) | |

| Microbiological taxa, n (%) | France N=819 | France, overseas territories[a] N=78 | Europe, other countries N=121 | North Africa N=389 | Sub-Saharan Africa N=264 | Asia N=64 | *P* value[c] |
|---|---|---|---|---|---|---|---|
| Rare | 24 (2.9) | 5 (6.4) | 4 (3.3) | 5 (1.3) | 12 (4.5) | 0 (0) | |
| Absent | 702 (85.9) | 67 (85.9) | 104 (86.0) | 333 (85.6) | 211 (79.9) | 55 (85.9) | |
| *Missing* | *2 (0.2)* | *0 (0)* | *0 (0)* | *0 (0)* | *0 (0)* | *0 (0)* | |
| ***Streptococcus anginosus*** | | | | | | | 0.43 |
| Numerous | 23 (2.8) | 2 (2.6) | 4 (3.3) | 11 (2.8) | 11 (4.2) | 3 (4.7) | |
| Rare | 10 (1.2) | 1 (1.3) | 3 (2.5) | 1 (0.3) | 1 (0.4) | 1 (1.6) | |
| Absent | 786 (96.0) | 75 (96.2) | 114 (94.2) | 376 (96.9) | 251 (95.4) | 60 (93.8) | |
| *Missing* | *0 (0)* | *0 (0)* | *0 (0)* | *1 (0.3)* | *1 (0.4)* | *0 (0)* | |
| **Anaerobes[b]** | | | | | | | <0.0001 |
| Numerous | 82 (10.0) | 27 (34.6) | 17 (14.0) | 33 (8.5) | 49 (18.6) | 8 (12.5) | |
| Rare | 12 (1.5) | 0 (0) | 2 (1.7) | 1 (0.3) | 7 (2.7) | 0 (0) | |
| Absent | 724 (88.5) | 51 (65.4) | 102 (84.3) | 354 (91.2) | 208 (78.8) | 56 (87.5) | |
| *Missing* | *1 (0.1)* | *0 (0)* | *0 (0)* | *1 (0.3)* | *0 (0)* | *0 (0)* | |
| ***Gardnerella* spp.** | | | | | | | <0.0001 |
| Numerous | 53 (6.5) | 22 (28.2) | 13 (10.7) | 28 (7.2) | 39 (14.8) | 6 (9.4) | |
| Rare | 8 (1.0) | 0 (0) | 2 (1.7) | 1 (0.3) | 5 (1.9) | 0 (0) | |
| Absent | 757 (92.5) | 56 (71.8) | 106 (87.6) | 359 (92.5) | 220 (83.3) | 58 (90.6) | |
| *Missing* | *1 (0.1)* | *0 (0)* | *0 (0)* | *1 (0.3)* | *0 (0)* | *0 (0)* | |
| ***Candida* spp.** | | | | | | | 0.0002 |
| Numerous | 93 (11.4) | 13 (16.9) | 18 (14.9) | 60 (15.5) | 62 (23.6) | 6 (9.4) | |
| Rare | 21 (2.6) | 3 (3.9) | 1 (0.8) | 10 (2.6) | 12 (4.6) | 0 (0) | |
| Absent | 699 (86.0) | 61 (79.2) | 102 (84.3) | 318 (82.0) | 189 (71.9) | 58 (90.6) | |
| *Missing* | *6 (0.7)* | *1 (1.3)* | *0 (0)* | *1 (0.3)* | *1 (0.4)* | *0 (0)* | |

[a] Including territories in South America (French Guiana), the Atlantic Ocean (Guadeloupe, Martinique, Saint-Martin, Saint-Barthélemy, Saint-Pierre-et-Miquelon), the Indian Ocean (Reunion Island, Mayotte), the Pacific Ocean (New Caledonia, French Polynesia, the Wallis and Futuna Islands), and the French Southern and Antarctic Territories.
[b] including *Gardnerella* spp.
[c] Chi-square tests.

**Supplementary Table 6.** Nugent score at enrollment according to gravidity and parity.

| | Gravidity | | | | Parity | | | |
|---|---|---|---|---|---|---|---|---|
| **Nugent score, n (%)** | **1**<br>**N=564** | **2**<br>**N=550** | **≥ 3**<br>**N=734** | ***P* value**[a] | **0**<br>**N=880** | **1**<br>**N=609** | **≥ 2**<br>**N=359** | ***P* value**[a] |
| Normal (0-3) | 415 (87.5) | 363 (83.9) | 430 (81.6) | 0.010 | 640 (86.4) | 372 (82.1) | 196 (81.3) | 0.041 |
| Intermediate (4-6) | 53 (11.2) | 56 (12.9) | 81 (15.4) | | 84 (11.3) | 72 (15.9) | 34 (14.1) | |
| Bacterial vaginosis (≥7) | 7 (1.5) | 14 (3.2) | 16 (3.0) | | 17 (2.3) | 9 (2.0) | 11 (4.6) | |
| *Missing* | *89 (15.8)* | *117 (21.3)* | *207 (28.2)* | | *139 (15.8)* | *156 (25.6)* | *118 (32.9)* | |

[a] Chi-square tests.

**Supplementary Table 7.** Microbiological cultures of vaginal samples at enrollment according to gravidity and parity.

| | Gravidity | | | | Parity | | | |
|---|---|---|---|---|---|---|---|---|
| **Microbiological taxa, n (%)** | **1 N=564** | **2 N=550** | **≥ 3 N=734** | ***P* value[b]** | **0 N=880** | **1 N=609** | **≥ 2 N=359** | ***P* value[b]** |
| ***Lactobacillus* spp.** | | | | <0.0001 | | | | 0.0001 |
| Numerous | 445 (80.2) | 396 (72.7) | 486 (66.7) | | 670 (77.4) | 423 (69.8) | 234 (65.7) | |
| Rare | 20 (3.6) | 29 (5.3) | 34 (4.7) | | 35 (4.0) | 33 (5.4) | 15 (4.2) | |
| Absent | 90 (16.2) | 120 (22.0) | 208 (28.6) | | 161 (18.6) | 150 (24.8) | 107 (30.1) | |
| *Missing* | *9 (1.6)* | *5 (0.9)* | *6 (0.8)* | | *14 (1.6)* | *3 (0.5)* | *3 (0.8)* | |
| **Enterobacteria, all species** | | | | 0.88 | | | | 0.74 |
| Numerous | 77 (13.6) | 75 (13.6) | 91 (12.4) | | 117 (13.3) | 76 (12.5) | 50 (14.0) | |
| Rare | 58 (10.3) | 49 (8.9) | 71 (9.7) | | 91 (10.3) | 52 (8.5) | 35 (9.8) | |
| Absent | 429 (76.1) | 426 (77.5) | 571 (77.9) | | 672 (76.4) | 481 (79.0) | 273 (76.2) | |
| *Missing* | *0* | *0* | *1 (0.1)* | | *0* | *0* | *1 (0.3)* | |
| ***Escherichia coli*** | | | | 0.59 | | | | 0.29 |
| Numerous | 56 (10.0) | 51 (9.3) | 55 (7.5) | | 86 (9.8) | 45 (7.4) | 31 (8.7) | |
| Rare | 45 (8.0) | 41 (7.5) | 55 (7.5) | | 75 (8.6) | 41 (6.7) | 25 (7.0) | |
| Absent | 461 (82.0) | 458 (83.2) | 621 (85.0) | | 716 (81.6) | 523 (85.9) | 301 (84.3) | |
| *Missing* | *2 (0.4)* | *0* | *3 (0.4)* | | *3 (0.3)* | *0* | *2 (0.6)* | |
| ***Klebsiella pneumoniae*** | | | | 0.83 | | | | 0.29 |
| Numerous | 14 (2.5) | 16 (2.9) | 26 (3.6) | | 22 (2.5) | 18 (3.0) | 16 (4.5) | |
| Rare | 9 (1.6) | 7 (1.3) | 10 (1.3) | | 13 (1.5) | 6 (1.0) | 7 (2.0) | |
| Absent | 538 (95.9) | 527 (95.8) | 696 (95.1) | | 841 (96.0) | 585 (96.0) | 335 (93.5) | |
| *Missing* | *3 (0.5)* | *0* | *2 (0.3)* | | *4 (0.5)* | *0* | *1 (0.3)* | |
| ***Staphylococcus aureus*** | | | | 0.13 | | | | 0.21 |
| Numerous | 13 (2.3) | 10 (1.8) | 29 (4.0) | | 23 (2.6) | 14 (2.3) | 15 (4.2) | |
| Rare | 11 (2.0) | 11 (2.0) | 19 (2.6) | | 17 (2.0) | 12 (2.0) | 12 (3.4) | |
| Absent | 532 (95.7) | 528 (96.2) | 675 (93.4) | | 828 (95.4) | 578 (95.7) | 329 (92.4) | |
| *Missing* | *8 (1.4)* | *1 (0.2)* | *11 (1.5)* | | *12 (1.4)* | *5 (0.8)* | *3 (0.8)* | |
| **Group B *Streptococcus*** | | | | 0.062 | | | | 0.046 |
| Numerous | 52 (9.3) | 74 (13.5) | 94 (12.8) | | 86 (9.8) | 77 (12.6) | 57 (15.9) | |

| | Gravidity | | | | Parity | | | |
|---|---|---|---|---|---|---|---|---|
| **Microbiological taxa, n (%)** | **1 N=564** | **2 N=550** | **≥ 3 N=734** | ***P* value[b]** | **0 N=880** | **1 N=609** | **≥ 2 N=359** | ***P* value[b]** |
| Rare | 21 (3.7) | 10 (1.8) | 23 (3.1) | | 28 (3.2) | 16 (2.6) | 10 (2.8) | |
| Absent | 489 (87.0) | 466 (84.7) | 617 (84.1) | | 764 (87.0) | 516 (84.8) | 292 (81.3) | |
| *Missing* | *2 (0.4)* | *0* | *0* | | *2 (0.2)* | *0* | *0* | |
| ***Streptococcus anginosus*** | | | | 0.082 | | | | 0.0006 |
| Numerous | 21 (3.7) | 13 (2.4) | 24 (3.3) | | 40 (4.5) | 8 (1.3) | 10 (2.8) | |
| Rare | 9 (1.6) | 7 (1.3) | 2 (0.3) | | 14 (1.6) | 3 (0.5) | 1 (0.3) | |
| Absent | 534 (94.7) | 529 (96.3) | 708 (96.4) | | 825 (93.9) | 598 (98.2) | 348 (96.9) | |
| *Missing* | *0* | *1 (0.2)* | *0* | | *1 (0.1)* | *0* | *0* | |
| **Anaerobes[a]** | | | | 0.23 | | | | 0.95 |
| Numerous | 64 (11.4) | 65 (11.8) | 108 (14.7) | | 113 (12.8) | 81 (13.3) | 43 (12.0) | |
| Rare | 4 (0.7) | 8 (1.5) | 10 (1.4) | | 11 (1.3) | 6 (1.0) | 5 (1.4) | |
| Absent | 495 (87.9) | 476 (86.7) | 615 (83.9) | | 755 (85.9) | 521 (85.7) | 310 (86.6) | |
| *Missing* | *1 (0.2)* | *1 (0.2)* | *1 (0.1)* | | *1 (0.1)* | *1 (0.2)* | *1 (0.3)* | |
| ***Gardnerella* spp.** | | | | 0.13 | | | | 0.63 |
| Numerous | 41 (7.3) | 53 (9.7) | 84 (11.5) | | 80 (9.1) | 65 (10.7) | 33 (9.2) | |
| Rare | 4 (0.7) | 4 (0.7) | 8 (1.1) | | 9 (1.0) | 3 (0.5) | 4 (1.1) | |
| Absent | 518 (92.0) | 492 (89.6) | 641 (87.4) | | 790 (89.9) | 540 (88.8) | 321 (89.7) | |
| *Missing* | *1 (0.2)* | *1 (0.2)* | *1 (0.1)* | | *1 (0.1)* | *1 (0.2)* | *1 (0.3)* | |
| ***Candida* spp.** | | | | 0.016 | | | | 0.015 |
| Numerous | 60 (10.7) | 78 (14 .2) | 125 (17.1) | | 99 (11.3) | 102 (16.9) | 62 (17.4) | |
| Rare | 12 (2.2) | 17 (3.1) | 22 (3.0) | | 25 (2.9) | 16 (2.6) | 10 (2.8) | |
| Absent | 487 (87.1) | 454 (82.7) | 583 (79.9) | | 751 (85.8) | 488 (80.5) | 285 (79.8) | |
| *Missing* | *5 (0.9)* | *1 (0.2)* | *4 (0.5)* | | *5 (0.6)* | *3 (0.5)* | *2 (0.6)* | |

[a] including *Gardnerella* spp.
[b] Chi-square tests.

**Supplementary Table 8.** Nugent scores at enrollment according to the body mass index.[a]

| **Nugent score, n (%)** | **Underweight N=114** | **Normal N=1,077** | **Overweight N=403** | **Obese N=242** | ***P* value[b]** |
|---|---|---|---|---|---|
| Normal (0-3) | 80 (76.9) | 751 (85.6) | 236 (83.1) | 131 (82.4) | 0.024 |
| Intermediate (4-6) | 21 (20.2) | 111 (12.7) | 38 (13.4) | 19 (11.9) | |
| Bacterial vaginosis (≥7) | 3 (2.9) | 15 (1.7) | 10 (3.5) | 9 (5.7) | |
| *Missing* | *10 (8.8)* | *200 (18.6)* | *119 (29.5)* | *83 (34.3)* | |

[a] Underweight: BMI < 18.5; Normal weight: BMI ≥ 18.5 and < 25; Overweight: BMI ≥ 25 and < 30; Obese: BMI ≥ 30.
[b] Chi-square test.

**Supplementary Table 9.** Microbiological cultures of vaginal samples at enrolment according to the body mass index.[a]

| Microbiological taxa, n (%) | Underweight N=114 | Normal N=1,077 | Overweight N=403 | Obese N=242 | *P* value[c] |
|---|---|---|---|---|---|
| ***Lactobacillus* spp.** | | | | | 0.0029 |
| Numerous | 89 (79.5) | 794 (74.6) | 289 (72.3) | 149 (61.8) | |
| Rare | 3 (2.7) | 45 (4.2) | 16 (4.0) | 17 (7.1) | |
| Absent | 20 (17.9) | 225 (21.1) | 95 (23.7) | 75 (31.1) | |
| *Missing* | *2 (1.8)* | *13 (1.2)* | *3 (0.7)* | *1 (0.4)* | |
| **Enterobacteria, all species** | | | | | <0.0001 |
| Numerous | 18 (15.8) | 114 (10.6) | 64 (15.9) | 45 (18.6) | |
| Rare | 8 (7.0) | 90 (8.4) | 44 (10.9) | 35 (14.5) | |
| Absent | 88 (77.2) | 873 (81.1) | 295 (73.2) | 162 (66.9) | |
| *Missing* | *0 (0)* | *0 (0)* | *0 (0)* | *0 (0)* | |
| ***Escherichia coli*** | | | | | 0.035 |
| Numerous | 13 (11.5) | 83 (7.7) | 38 (9.4) | 26 (10.7) | |
| Rare | 5 (4.4) | 70 (6.5) | 40 (9.9) | 25 (10.3) | |
| Absent | 95 (84.1) | 921 (85.8) | 325 (80.6) | 191 (78.9) | |
| *Missing* | *1 (0.9)* | *3 (0.3)* | *0 (0)* | *0 (0)* | |
| ***Klebsiella pneumoniae*** | | | | | 0.0002 |
| Numerous | 2 (1.8) | 19 (1.8) | 18 (4.5) | 17 (7.0) | |
| Rare | 1 (0.9) | 11 (1.0) | 8 (2.0) | 6 (2.5) | |
| Absent | 110 (97.3) | 1043 (97.2) | 377 (93.5) | 219 (90.5) | |
| *Missing* | *1 (0.9)* | *4 (0.4)* | *0 (0)* | *0 (0)* | |
| ***Staphylococcus aureus*** | | | | | 0.74 |
| Numerous | 3 (2.7) | 32 (3.0) | 12 (3.0) | 5 (2.1) | |
| Rare | 3 (2.7) | 18 (1.7) | 10 (2.5) | 8 (3.3) | |
| Absent | 107 (94.7) | 1009 (95.3) | 380 (94.5) | 229 (94.6) | |
| *Missing* | *1 (0.9)* | *18 (1.7)* | *1 (0.2)* | *0 (0)* | |
| **Group B *Streptococcus*** | | | | | 0.84 |
| Numerous | 11 (9.6) | 128 (11.9) | 53 (13.1) | 27 (11.2) | |
| Rare | 5 (4.4) | 32 (3.0) | 12 (3.3) | 5 (2.1) | |
| Absent | 98 (86.0) | 915 (85.1) | 338 (83.7) | 210 (86.8) | |

| Microbiological taxa, n (%) | Underweight N=114 | Normal N=1,077 | Overweight N=403 | Obese N=242 | *P* value[c] |
|---|---|---|---|---|---|
| *Missing* | *0 (0)* | *2 (0.2)* | *0 (0)* | *0 (0)* | |
| ***Streptococcus anginosus*** | | | | | 0.098 |
| Numerous | 3 (2.6) | 28 (2.6) | 20 (5.0) | 7 (2.9) | |
| Rare | 1 (0.9) | 15 (1.4) | 2 (0.5) | 0 | |
| Absent | 110 (96.5) | 1033 (96.0) | 381 (94.5) | 235 (97.1) | |
| *Missing* | *0 (0)* | *1 (0.1)* | *0 (0)* | *0 (0)* | |
| **Anaerobes[b]** | | | | | 0.55 |
| Numerous | 18 (15.9) | 140 (13.0) | 49 (12.2) | 29 (12.0) | |
| Rare | 0 (0) | 17 (1.6) | 3 (0.7) | 2 (0.8) | |
| Absent | 95 (84.1) | 919 (85.4) | 350 (87.1) | 211 (87.2) | |
| *Missing* | *1 (0.9)* | *1 (0.1)* | *1 (0.2)* | *0 (0)* | |
| ***Gardnerella* spp.** | | | | | 0.49 |
| Numerous | 14 (12.4) | 103 (9.6) | 36 (9.0) | 24 (9.9) | |
| Rare | 0 (0) | 13 (1.2) | 1 (0.2) | 2 (0.8) | |
| Absent | 99 (87.6) | 960 (89.2) | 365 (90.8) | 216 (89.3) | |
| *Missing* | *1 (0.9)* | *1 (0.1)* | *1 (0.2)* | *0 (0)* | |
| ***Candida* spp.** | | | | | 0.029 |
| Numerous | 10 (8.8) | 157 (14.6) | 52 (13.0) | 42 (17.4) | |
| Rare | 1 (0.9) | 27 (2.5) | 10 (2.5) | 13 (5.4) | |
| Absent | 102 (90.3) | 889 (82.9) | 338 (84.5) | 186 (77.2) | |
| *Missing* | *1 (0.9)* | *4 (0.4)* | *3 (0.7)* | *1 (0.4)* | |

[a] Underweight: BMI < 18.5; Normal weight: BMI ≥ 18.5 and < 25; Overweight: BMI ≥ 25 and < 30; Obese: BMI ≥ 30.
[b] Including *Gardnerella* spp.
[c] Chi-square tests.
