## Supplemental Figures for "Culture-based analysis of vaginal microbiota and its link to pregnancy outcomes: an observational cohort study in France"

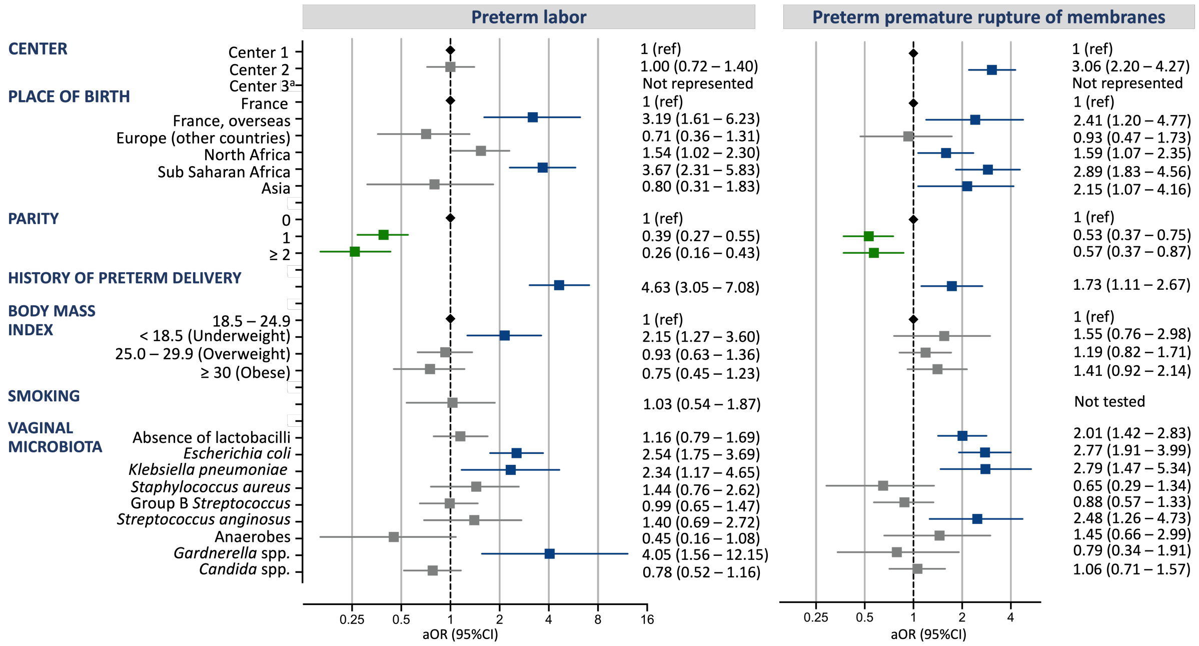


**Supplementary Figure 1. Second model of multivariable analysis of risk factors for preterm labor and preterm premature rupture of membranes.**

In this model, *Escherichia coli* and *Klebsiella pneumoniae* were used instead of enterobacteria as variables. Error bars indicate the upper and lower limit of the 95% confidence interval (CI); aOR: adjusted odds ratio. [a] aOR (95% CI) of Center 3 for preterm labor and preterm premature rupture of membranes: $1.6 \times 10^{8}$ ($6.7 \times 10^{2}$-$2.5 \times 10^{70}$) and $2.4 \times 10^{8}$ (0.09-$6.3 \times 10^{85}$), respectively.

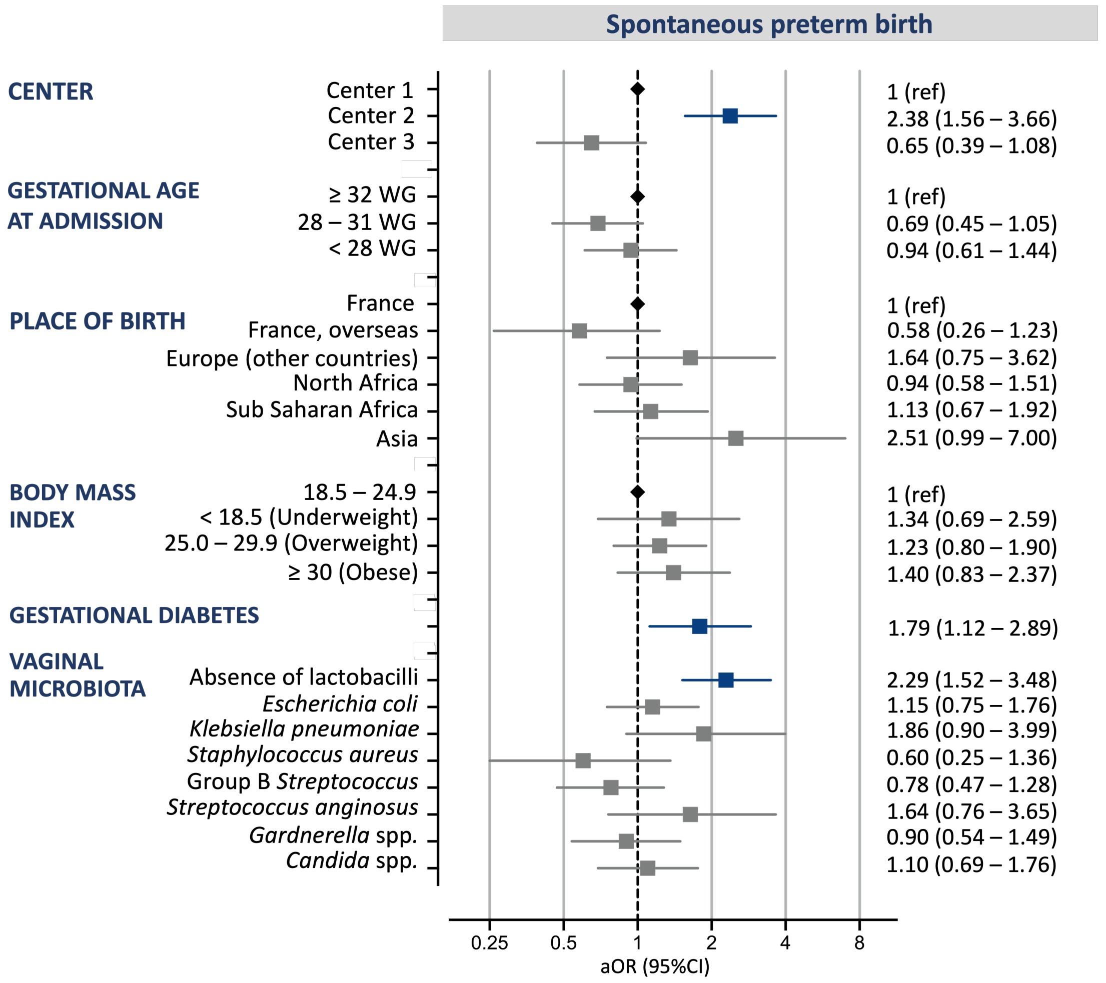


**Supplementary Figure 2. Second model of multivariable analysis of risk factors for spontaneous preterm birth.**

In this model, *Escherichia coli* and *Klebsiella pneumoniae* were used instead of enterobacteria as variables. Error bars indicate the upper and lower limit of the 95% confidence interval (CI); aOR: adjusted odds ratio. WG: weeks of gestation.
